# Inactivation of the mitochondrial protease Afg3l2 results in severely diminished respiratory chain activity and widespread defects in mitochondrial gene expression

**DOI:** 10.1101/2020.05.31.126607

**Authors:** Gautam Pareek, Leo J. Pallanck

**Affiliations:** Department of Genome Sciences, University of Washington, 3720 15^th^ Avenue NE, Seattle, WA 98195, USA

## Abstract

The *m*-AAA proteases plays a critical role in the proteostasis of the inner mitochondrial membrane proteins, and mutations in the genes encoding these proteases cause severe incurable neurological diseases. To further explore the biological role of the *m*-AAA proteases and the pathological consequences of their deficiency, we used a genetic approach in the fruit fly *Drosophila melanogaster* to inactivate the ATPase family gene 3-like 2 (*AFG3L2*) gene, which encodes a component of the *m*-AAA proteases. We found that null alleles of *Drosophila AFG3L2* die early in development, but partial inactivation of *AFG3L2* using RNAi extended viability to the late pupal and adult stages of development. Flies with partial inactivation of Afg3l2 exhibited marked behavioral defects, neurodegeneration, mitochondrial morphological alterations, and diminished respiratory chain (RC) activity. Further work revealed that reduced RC activity was a consequence of widespread defects in mitochondrial gene expression, including diminished mitochondrial transcription, translation and impaired mitochondrial ribosome biogenesis. These defects were accompanied by the compensatory activation of the mitochondrial unfolded protein response (mito-UPR) and accumulation of unfolded mitochondrial proteins, including proteins involved in transcription. Overexpression of the mito-UPR components partially rescued the Afg3l2-deficient phenotypes, indicating that sequestration of essential components of the mitochondrial gene expression into aggregates partly accounts for these defects. However, Afg3l2 also co-sediments with the mitochondrial ribosome biogenesis machinery, suggesting an additional novel role for Afg3l2 in ribosome biogenesis. Our work suggests that strategies designed to modify mitochondrial stress pathways and mitochondrial gene expression could be therapeutic in the diseases caused by mutations in *AFG3L2*.

**Author Summary:** Mitochondria produce virtually all of the cellular energy through the actions of the respiratory chain (RC) complexes. However, both the assembly of the RC complexes, and their biological functions come at a cost. Biogenesis of the RC complexes depends on the coordinated expression of nuclear and mitochondrially encoded subunits and an imbalance in this process can cause protein aggregation. Moreover, the RC complexes produce highly damaging reactive oxygen species as a side product of their activity. The Mitochondrial AAA^+^ family of proteases are believed to provide the first line of defense against these insults. The importance of this protease family is best exemplified by the severe neurodegenerative diseases that are caused by mutations in their respective genes. To better understand the biological roles of the AAA^+^ proteases, and the physiological consequences of their inactivation we used a genetic approach in *Drosophila* to study the Afg3l2 AAA^+^ protease. Unexpectedly, we found that Afg3l2 deficiency profoundly impaired mitochondrial gene expression, including transcription, translation and ribosome biogenesis. These phenotypes were accompanied by accumulation of insoluble mitochondrial proteins, and compensatory activation of mito-UPR and autophagy. Our work indicates Afg3l2 plays critical roles in degrading unfolded mitochondrial proteins and regulating mitochondrial gene expression.

## Introduction

Mitochondria play a number of indispensable roles, most notably the production of virtually all cellular energy by coupling the mitochondrial membrane potential created by the respiratory chain (RC) complexes to the synthesis of ATP (1). However, the RC complexes also produce reactive oxygen species (ROS) that can damage mitochondrial constituents (2). Moreover, proper assembly of the RC complexes requires the coordinated expression of mitochondrial and nuclear genome encoded RC subunits, and an imbalance in the stoichiometry of these subunits can cause protein misfolding and aggregation (3). To preserve mitochondrial integrity, eukaryotes have evolved a variety of quality control strategies, including in extreme circumstances the degradation of the mitochondria in the lysosome through a mitochondrial selective form of autophagy known as mitophagy (4) (5). The relatively recent demonstration that the Parkinson’s disease genes *PINK1* and *PARKIN* play critical roles in mitophagy has led to a broad interest in this mitochondrial quality control pathway. However, a number of observations suggest that mitophagy accounts for only a minority of mitochondrial protein turnover, indicating that other quality control processes predominate (6).

The AAA^+^ family of mitochondrial proteases represents likely candidates to account for the majority of mitochondrial protein turnover (7). This family of proteases forms oligomeric complexes that use energy from ATP hydrolysis to unfold and transport substrates to their proteolytic cavity for degradation. Higher eukaryotes possess four different mitochondrial AAA^+^ proteases that localize to different mitochondrial sub-compartments (8). One of these four proteases, designated *m-*AAA, is embedded in the inner membrane with its active site oriented towards the matrix. There are two different versions of *m-*AAA; one version is composed entirely of the Afg3l2 protein; the other version is a hetero-oligomeric complex of the Afg3l2 and Paraplegin proteins (9). Mutations in the genes encoding Afg3l2 and Paraplegin cause spinocerebellar ataxia (SCA28) and hereditary spastic paraplegia (HSP) respectively, but the interplay between these proteins, their substrates, and the pathways influenced by mutations in their respective genes are poorly understood (8).

To explore the biological functions of the AAA^+^ mitochondrial protease family, we are using a genetic approach in the fruit fly *Drosophila* (10–12). In our current work, we describe the phenotypes caused by the inactivation of *AFG3L2*. Null alleles of *AFG3L2* die early in larval development, but partial inactivation of *AFG3L2* using RNAi resulted in viable adult animals with a significantly shortened lifespan, locomotor impairment, and neurodegeneration. These phenotypes were accompanied by a wide variety of mitochondrial defects, including a fragmented morphology, reduced cristae density, an accumulation of insoluble mitochondrial proteins and a reduction in the activity and abundance of the RC complexes resulting from impaired mitochondrial gene expression. The defects in mitochondrial gene expression appear to derive in part from reduced transcription caused by sequestration of transcription factors into insoluble aggregates. However, the defect in proteolytic processing of the large mitochondrial ribosome subunit Mrpl32 and impaired assembly of the mitochondrial ribosomes also contributes to the translational defect of Afg3l2 deficiency. Together, our findings indicate that the pathological consequences of Afg3l2 deficiency derive from protein unfolding and diminished mitochondrial gene expression and that measures to reverse these defects would be therapeutic in the diseases caused by mutations in *AFG3L2*.

## Results

### *AFG3L2* is an essential gene in *Drosophila*

In previous work, we established that *CG6512* encodes the *Drosophila* Afg3l2 ortholog, in part because it exhibits greater than 60% amino acid identity with human Afg3l2 over most of its sequence (13). In further support of this matter, we confirmed the mitochondrial localization of the *CG6512* gene product using immunofluorescence microscopy (Supplemental Figure 1A) and subcellular fractionation (Supplemental Figure 1B).

To create a *Drosophila* model of the diseases caused by mutations in *AFG3L2*, we used the CRISPR/Cas9 system to replace the entire *Drosophila AFG3L2* coding sequence with the DsRed marker (14). Previous work has shown that heterozygous mutations in the human *AFG3L2* gene cause spinocerebellar ataxia type 28 (SCA28), and haploinsufficiency of *AFG3L2* in mice has been shown to cause mitochondria-mediated Purkinje cell dark degeneration (15, 16). However, flies bearing a single copy of our null allele of *AFG3L2* (designated *AFG3L2^del^*) were viable and fertile and exhibited no significant difference in lifespan relative to controls (median lifespan of *AFG3L2^del^*/+ flies = 64 days compared to 66 days for control flies), indicating that haploinsufficiency of *AFG3L2* is well tolerated in *Drosophila* (Figure 1A and Supplemental Table 1). Western blot analysis of the *AFG3L2^del^* heterozygotes confirmed the expected reduction in Afg3l2 protein abundance, indicating that the lack of phenotype in *AFG3L2^del^* heterozygotes was not a consequence of compensatory upregulation of Afg3l2 protein expression in the heterozygotes (Figure 1B). By contrast, *AFG3L2^del^* homozygotes died before the end of the second instar larval stage of development, indicating that the *AFG3L2* gene is essential for viability (Figure 1C). The recessive lethal phenotype conferred by the *AFG3L2^del^* allele was rescued by ectopic expression of *AFG3L2* using the ubiquitous Daughterless (da) – Gal4 driver, verifying that this phenotype is due to *AFG3L2* inactivation (Supplemental Figure 2A-B).

**Figure 1.**
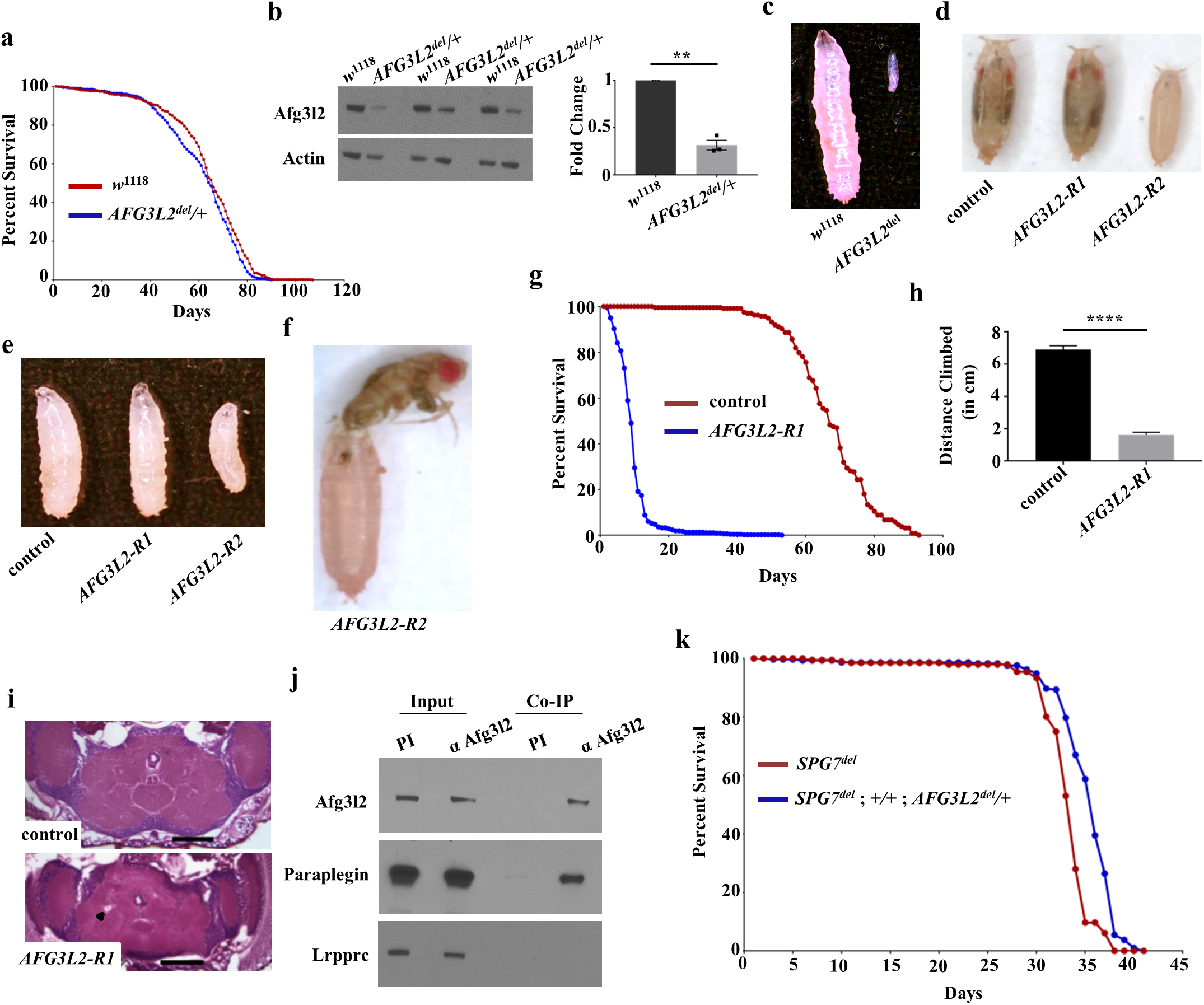
*AFG3L2* null mutants die early in development, but neuron-specific inactivation of *AFG3L2* confers shortened adult lifespan, locomotor defects, and neurodegeneration. **a.** Kaplan–Meier survival curves of control (*w*^1118^; N = 480, 50% survival 66 days) and *AFG3L2* heterozygotes (*AFG3L2^del^*/+; N = 548, 50% survival 65 days). **b.** Western blot analysis of protein homogenates from 1-day old controls (*w*^1118^) and *AFG3L2* heterozygotes (*AFG3L2^del^***/+**) using antisera against Afg3l2 and actin. Afg3l2 band intensity was normalized against actin (n = 3 independent biological replicates, **p < 0.01 by Student’s t-test). **c.** Size comparison of control (*w*^1118^) and *AFG3L2* null homozygous larvae (*AFG3L2^del/del^*) 4 days after egg hatching. **d, e.** Size comparison of control animals (UAS-*LUCIFERASE* RNAi/da-Gal4) and flies expressing RNAi lines targeting *AFG3L2* in a ubiquitous manner (UAS-*AFG3L2-R1* RNAi; da-Gal4 or UAS-*AFG3L2-R2* RNAi/da-Gal4). Images show pupae 9 days after egg hatching **(d)** and larvae 4 days after egg hatching **(e)**. **f.** Pan-neuronal expression of the strong RNAi line targeting *AFG3L2* (elav-Gal4; +/+; UAS-*AFG3L2-R2* RNAi) resulted in the failure of flies to eclose from the pupal case. **g.** Kaplan–Meier survival curves of flies expressing the control *LUCIFERASE* RNAi (elav-Gal4; +/+; UAS-*LUCIFERASE* RNAi; N = 238, 50% survival 67 days) and weak RNAi line targeting *AFG3L2* (elav-Gal4; UAS-*AFG3L2-R1* RNAi; N = 465, 50% survival 9 days) throughout the nervous system. Significance was determined using a Mantel-Cox log-rank test (\*\*\*\**p* < 0.0001). **h.** Climbing performance of 1-day old adult flies expressing the control *LUCIFERASE* RNAi (elav-Gal4; +/+; UAS-*LUCIFERASE* RNAi; N = 70) and the weak RNAi line targeting *AFG3L2* (elav-Gal4; UAS-*AFG3L2-R1* RNAi; N = 75) throughout the nervous system (\*\*\*\**p* < 0.0005 by Student’s t-test). **i.** Hematoxylin and eosin staining of paraffin-embedded head sections from 1-day old controls (elav-Gal4; +/+; UAS-*LUCIFERASE* RNAi) and flies expressing the weak RNAi line targeting *AFG3L2* (elav-Gal4; UAS-*AFG3L2-R1* RNAi) throughout the nervous system. The black arrowhead indicates a vacuole. The scale bar is 100 μm. **j.** Western blot analysis of *Drosophila* proteins immunoprecipitated using an antiserum against Afg3l2 (α Afg3l2). A pre-immune serum (PI) was used as a negative control. The blot was incubated with antiserum against Afg3l2 to confirm successful immunoprecipitation of Afg3l2, and an antiserum against Paraplegin to test for co-immunoprecipitation of Paraplegin. An antiserum against Leucine-Rich Pentatricopeptide Repeat Containing (LRPPRC) protein was used as a negative control. **k.** Kaplan–Meier survival curves of Paraplegin null mutants (*SPG7^del^*; N = 196, 50% survival 34 days) and Paraplegin null mutants that also bear a null allele of *AFG3L2* (*SPG7^del^*; *+/+*; *AFG3L2^del^/+*; N = 291, 50% survival 36 days).

### Partial inactivation of *AFG3L2* causes shortened lifespan, locomotor deficits, and neurodegeneration

To identify conditions of *AFG3L2* inactivation that confer phenotypes that lie between the severe early larval lethality of *AFG3L2^del^* homozygotes and the phenotypically normal *AFG3L2^del^* heterozygotes, we tested several different RNAi lines targeting *AFG3L2* (13). We found that driving two different RNAi transgenes targeting the *AFG3L2* transcript (designated as *AFG3L2-R1* and *AFG3L2-R2*) using the da-Gal4 driver allowed survival to the pharate adult stage of development (Figure 1D). The *AFG3L2-R2* line appeared to be stronger than the *AFG3L2-R1* line, as indicated by the smaller sized larvae and pupae resulting from expression of this transgene relative to *AFG3L2-R1* (Figure 1D and E; Supplemental Figure 3 A and B). We further examined the phenotypes conferred by these RNAi lines using the pan-neuronal elav-Gal4 driver. Driving the stronger *AFG3L2-R2* transgene with the elav-Gal4 driver again resulted largely in pupal lethality with occasional adult escapers (Figure 1F). However, driving the weaker *AFG3L2-R1* transgene with the elav-Gal4 driver resulted in viable adults with no obvious morphological alterations upon eclosion. However, these flies were significantly shorter-lived than controls (median lifespan of *AFG3L2-R1* flies = 9 days compared to 67 days for control flies) (Figure 1G and Supplemental Table 1) and also exhibited severe locomotor deficits (Figure 1H). Histological analysis of paraffin-embedded brain sections from flies bearing the *AFG3L2-R1* and elav-Gal4 transgenes also revealed an increase in the number of brain vacuoles relative to age-matched controls (Figure 1I and Supplemental Figure 4), indicating that these flies have a neurodegenerative phenotype that may model the neurodegeneration characteristic of the diseases associated with mutations in *AFG3L2*.

### Afg3l2 assembles with Paraplegin in *Drosophila*, but haploinsufficiency of *AFG3L2* does not influence the phenotypes of Paraplegin deficiency

Vertebrates produce two distinct *m-*AAA proteases, one composed entirely of Afg3l2 and a second composed of Afg3l2 and Paraplegin, which is encoded by the *SPG7* gene in humans (17). It has also been shown that haploinsufficiency of *AFG3L2* exacerbates the axonopathy observed in Paraplegin-deficient mice, thus supporting the functional relationship between these two proteases (18). To test whether Afg3l2 and Paraplegin also assemble into a functional heteromultimeric complex in *Drosophila*, we performed two experiments. First, we tested whether these proteins physically interact with one another. We found that Afg3l2 can coimmunoprecipitate Paraplegin, whereas no such co-immunoprecipitation was observed with a control protein that does not interact with Afg3l2 (Figure 1J). This finding strongly suggests that *Drosophila* also forms an *m-*AAA protease consisting of a heteromultimeric complex of Afg3l2 and Paraplegin. Second, we tested whether haploinsufficiency of *AFG3L2* influences the lifespan of a *Drosophila SPG7* null mutant, designated *SPG7^del^*. In previously published work, we found that *SPG7^del^* mutants live only half as long as control flies (11). However, introduction of the *AFG3L2^del^* allele into this *SPG7* null background did not further shorten the lifespan of *SPG7^del^* mutants (median lifespan of *SPG7^del^; AFG3L2^del^*/+ = 34 days compared to 32 days for *SPG7^del^* flies) (Figure 1K and Supplemental Table 1). Together, our findings indicate that Afg3l2 and Paraplegin form a multimeric complex, but that loss of this complex does not confer sensitivity to Afg3l2 dosage alterations.

### Afg3l2 deficiency affects mitochondrial morphology

To begin to explore the impact of Afg3l2 deficiency on mitochondrial integrity, we analyzed mitochondrial ultrastructure in Afg3l2-deficient animals using transmission electron microscopy (19). Because the indirect flight muscles harbor numerous large mitochondria of relatively uniform size and cristae density, we performed this analysis using this tissue type by driving the *AFG3L2-R2* RNAi transgene in muscle tissue using the mesoderm-specific 24B-Gal4 driver (20). Animals expressing this transgene from the 24B-Gal4 driver died at the late pupal stage of development, so we performed our study using indirect flight muscles dissected from pupae. This analysis revealed that mitochondria in Afg3l2-deficient animals exhibited widespread reduction or loss of cristae density relative to controls (Figure 2A-B). Similar defects in mitochondrial ultrastructure have been reported in other AAA^+^ protease mutants (10, 11, 21).

**Figure 2.**
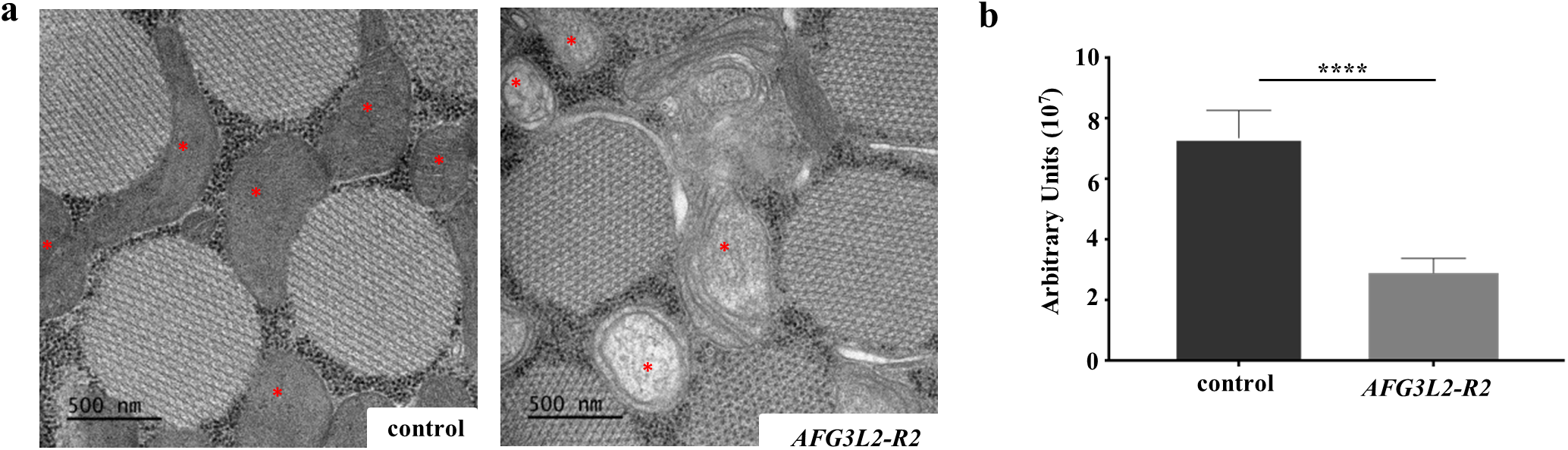
Afg3l2 deficiency results in mitochondrial morphological defects. **a.** Transmission electron microscopy images of indirect flight muscles from control animals (UAS-*LUCIFERASE* RNAi/24B-Gal4) and from flies expressing the strong *AFG3L2-R2* RNAi (UAS-*AFG3L2-R2* RNAi/24B-Gal4) using the mesoderm specific 24B-Gal4 driver. Images are of indirect flight muscles dissected from pupae. The red asterisks denote the mitochondria. **b.** Quantification of mitochondrial cristae density in control (UAS-*LUCIFERASE* RNAi/24B-Gal4) and Afg3l2-deficient (UAS-*AFG3L2-R2* RNAi/24B-Gal4) animals (\*\*\*\**p* < 0.0005 by Student’s t-test).

### Afg3l2 depletion impairs RC activity but does not influence mitochondrial calcium uptake kinetics

To examine the influence of Afg3l2 deficiency on mitochondrial function, we assayed two fundamental mitochondrial activities: respiratory chain activity and calcium uptake kinetics. Previous work has established that deficiency of the mitochondrial proteases Lon, dYmell1, and Paraplegin all result in both reduced activity and abundance of the RC complexes (10–12). To test whether similar RC defects occur upon depletion of Afg3l2, we measured RC activity in larvae and pupae bearing either the *AFG3L2-R1* or *AFG3L2-R2* transgenes along with a da-Gal4 driver. We found that the activity of complex I, III and IV were significantly reduced by Afg3l2 deficiency at both stages of development, although the magnitude of the reduction was greater in pupae (Figure 3A) than in larvae (Supplemental Figure 5). By contrast, complex II activity was unaffected (Figure 3A and Supplemental Figure 5). Consistent with these RC activity defects, we also observed a reduction in total ATP content in Afg3l2-deficient animals (Figure 3B). Because RC impairment is often accompanied by increased ROS production, we also compared total protein carbonylation content between Afg3l2-deficient animals and controls, an indicator of increased oxidative stress (22). However, our analysis revealed no apparent differences in this parameter between Afg3l2-deficient animals and controls (Supplemental Figure 6).

**Figure 3.**
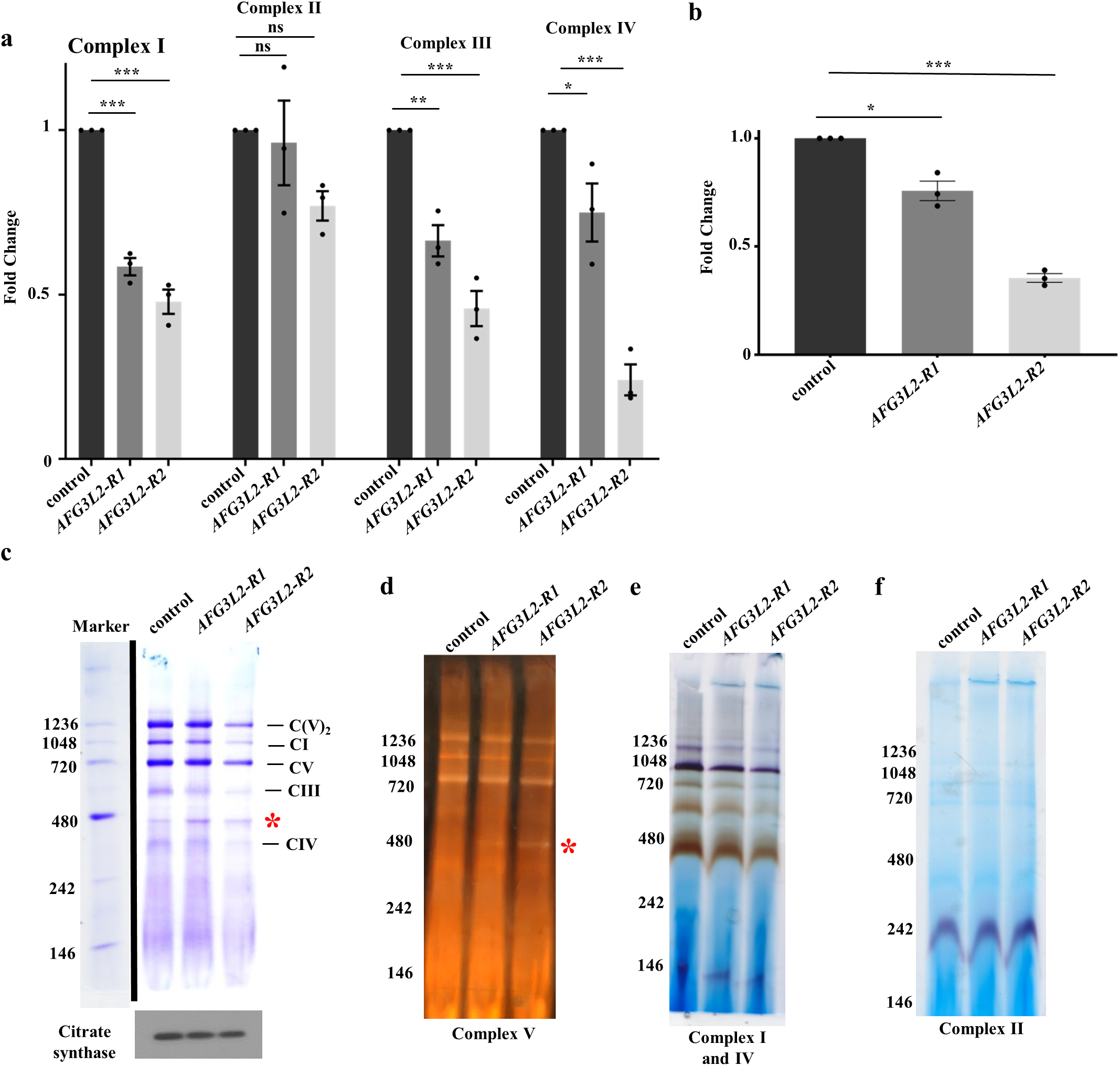
Afg3l2 deficiency confers diminished RC activity and abundance. **a**. RC activity in mitochondria isolated from control (UAS-*LUCIFERASE* RNAi/da-Gal4) and Afg3l2-deficient (UAS-*AFG3L2-R1* RNAi; da-Gal4 and UAS-*AFG3L2-R2* RNAi/da-Gal4) pupae (N = 3 independent biological replicates, \**p* < 0.05, \*\**p* < 0.005, \*\*\**p* < 0.0005 by one way ANOVA Tukey’s test for multiple comparison). **b.** ATP abundance in control (UAS-*LUCIFERASE* RNAi/da-Gal4) and Afg3l2-deficient (*UAS-AFG3L2-R1* RNAi; da-Gal4 and UAS-*AFG3L2-R2* RNAi/da-Gal4) pupae (N = 3 independent groups of 5 pupae, \**p* < 0.05, \*\*\**p* < 0.0005 by one way ANOVA Tukey’s test for multiple comparison). **c.** BN-PAGE of mitochondrial protein extracts from control (UAS-*LUCIFERASE* RNAi/da-Gal4) and Afg3l2-deficient (UAS-*AFG3L2 R1* RNAi; da-Gal4 and UAS-*AFG3L2-R2* RNAi/da-Gal4) pupae. Citrate synthase abundance was used as a loading control. Results of in-gel activity assays of mitochondrial complexes V **(d)**, I and IV **(e)**, and II **(f)** using the mitochondrial protein extracts from **(c)**. The red asterisk in **(c)** and **(d)** denotes the location of the sub-complex containing the F1 subunit of ATP synthase in Afg3l2-deficient animals. For BN-PAGE and in-gel activity assays, images shown here are representative of two independent biological replicates.

The decreases in complex I, III and IV activity in Afg3l2-deficient animals could result from functional defects in these complexes or a reduction in their abundance. To distinguish these possibilities, we used a combination of Blue Native PAGE (BN-PAGE), in-gel enzyme activity assays, and western blotting of selected subunits of these complexes to measure their abundance and to assess the efficiency of their assembly in Afg3l2-deficient animals. The use of these assay systems revealed that the abundance of all three of these complexes (Figure 3C-F), as well as their subunits, was reduced in Afg3l2-deficient animals (Supplemental Figure 7). We also detected the appearance of a catalytically active F1 subunit-containing sub-complex of ATP synthase (complex V) in Afg3l2-deficient animals, indicating that complex V assembly was also impaired (Figure 3D and Supplemental Figure 8). This assembly defect was also accompanied by an increase in the F1 subunit component ATPsynbeta (ATPVβ), supporting the idea that this assembly intermediate accumulates in Afg3l2-deficient animals (Supplemental Figure 7).

We next examined whether Afg3l2 deficiency influences mitochondrial calcium dynamics. The mitochondrial calcium uniporter (MCU) plays an important role in this process and recent reports indicate that the *m-AAA* proteases directly regulate MCU activity by degrading a positive regulatory subunit of this complex, known as the Essential MCU REgulator (EMRE) (23, 24). Degradation of free unassembled EMRE by the *m-*AAA proteases reduces the formation of a constitutively active MCU-EMRE subcomplex, thereby preventing calcium overload-triggered opening of the mitochondrial permeability transition pore (PTP) and cell death (25). To explore the potential influence of Afg3l2 deficiency on mitochondrial calcium dynamics, we first tested whether Afg3l2 regulates the abundance of EMRE or other MCU subunits by co-expressing transgenes encoding tagged forms of these proteins, including *EMRE*, *MICU1*, and *MCU* along with the *AFG3L2* RNAi transgenes using the da-Gal4 driver (26). We observed a significant increase in the abundance of Emre protein upon knockdown of Afg3l2, consistent with the conclusion that Afg3l2 is a negative regulator of Emre (Supplemental Figure 9A). However, we also observed similar increases in the abundance of Micu1 (Supplemental Figure 9B) and Mcu (Supplemental Figure 9C) upon knockdown of *AFG3L2*, suggesting that Afg3l2 plays a broader role in regulating the calcium uniporter than previously recognized.

Our finding that the abundance of all tested components of the calcium uniporter was increased upon deficiency of Afg3l2 suggested that the calcium uptake properties of mitochondria might also be altered in these animals and that these alterations might underlie the phenotypes of Afg3l2-deficient animals. However, because calcium uptake requires an intact membrane potential across the mitochondrial inner membrane (27, 28), we first analyzed if knockdown of Afg3l2 influences the mitochondrial membrane potential. Surprisingly, despite the mitochondrial morphological and functional deficits documented in Afg3l2-deficient animals, we detected no impairment in mitochondrial membrane potential upon expression of either the weak or strong *AFG3L2* RNAi lines in pupae using a da-Gal4 driver (Figure 4A). Thus, we proceeded to measure the calcium uptake properties of mitochondria from Afg3l2-deficient animals using established procedures (29). These experiments failed to detect any differences in the calcium uptake properties of mitochondria from Afg3l2-deficient animals relative to controls (Figure 4B). Nevertheless, we tested whether an RNAi targeting the core Ca^2+^ channel subunit MCU (designated as *MCU-R*) could rescue the pupal lethal phenotype of animals co-expressing the *AFG3L2* RNAi constructs using the da-Gal4 driver. In agreement with our finding that Afg3l2 deficiency did not impair mitochondrial calcium import, we found that knockdown of *MCU-R* failed to suppress the pupal lethal phenotype of animals expressing the *AFG3L2* RNAi construct from the da-Gal4 driver. Together, our findings indicate that Afg3l2 does not influence MCU activity and that mitochondrial calcium overload is not a feature of the pathogenic mechanism associated with Afg3l2 deficiency (Figure 4C).

**Figure 4.**
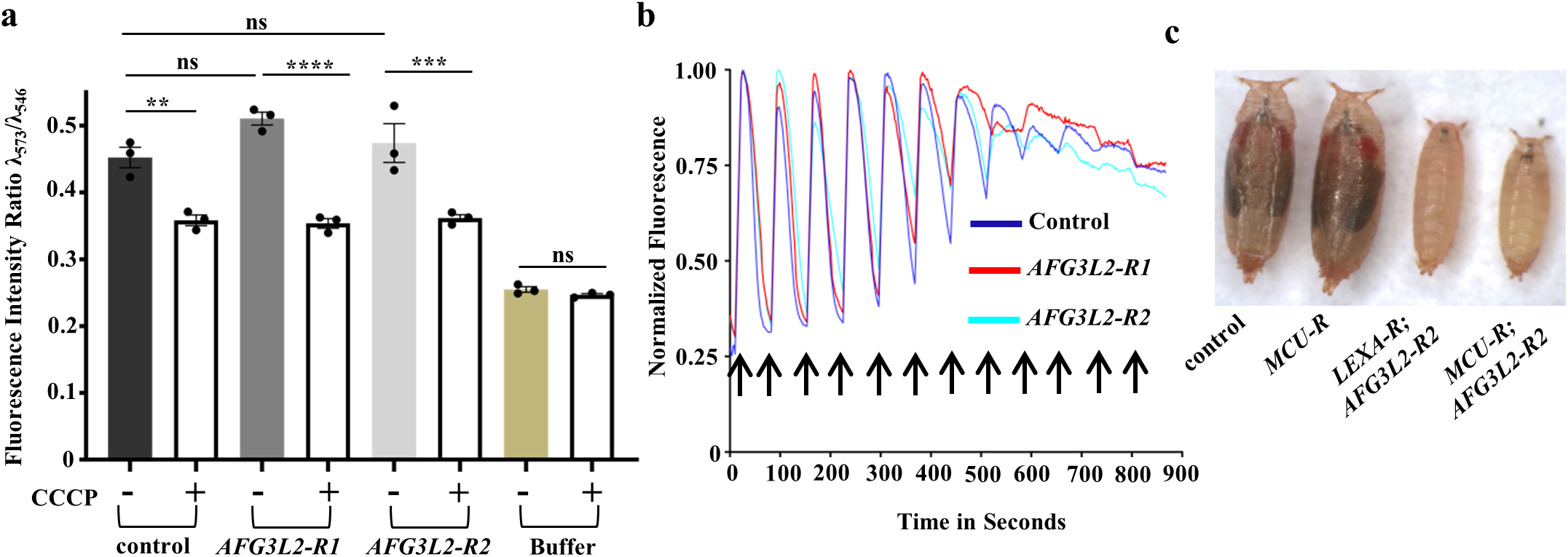
Mitochondrial membrane potential and calcium uptake are unaffected in Afg3l2-deficient animals. **a.** The mitochondrial membrane potential was examined using mitochondria from control (UAS-*LUCIFERASE* RNAi/da-Gal4) and Afg3l2-deficient (UAS-*AFG3L2-R1* RNAi; da-Gal4 and UAS-*AFG3L2-R2* RNAi/da-Gal4) pupae. Relative membrane potential was determined by measuring the fluorescence intensity ratio of Tetramethylrhodamine methyl ester perchlorate at 590nm using excitation wavelengths of 546 and 573nm (see materials and methods for details). CCCP (0.5 mM) was added to samples following the fluorescence scan to normalize for background fluorescence (\*\**p* < 0.01, \*\*\**p* < 0.0005, \*\*\*\**p* < 0.0001 by one way ANOVA Tukey’s test for multiple comparison). **b.** Calcium uptake capacity of mitochondria isolated from control (UAS-*LUCIFERASE* RNAi/da-Gal4) and Afg3l2-deficient (UAS-*AFG3L2-R1* RNAi; da-Gal4 and UAS-*AFG3L2-R2* RNAi/da-Gal4) pupae were determined using fluorescent dye Oregon green BAPTA 6F. The arrows represent the pulses of 500 μM CaCl2 after every 1 minute. Experiments were performed using two independent biological replicates and a representative trace is shown. **c.** Size comparison of control pupae (UAS-*LUCIFERASE* RNAi/da-Gal4), pupae expressing an RNAi targeting MCU: *MCU-R* (UAS-*MCU* RNAi; da-Gal4), Afg3l2-deficient pupae: *LEXA-R; AFG3L2-R2* (UAS-*LEXA* RNAi; UAS-*AFG3L2-R2* RNAi/da-Gal4) and Afg3l2-deficient pupae expressing an RNAi targeting MCU: *MCU-R; AFG3L2-R2* (UAS-*MCU* RNAi; UAS-*AFG3L2-R2* RNAi/da-Gal4). To account for the possible titration of Gal4 protein in the presence of two UAS transgenes, we compared *MCU-R*; *AFG3L2-R*2 to *AFG3L2-R*2 expressing the RNAi against the exogenous *LEXA* sequence (*LEXA-R*). Pupae in images were collected 9 days after egg hatching.

### Afg3l2 deficiency impairs mitochondrial transcription and translation

The alterations in RC activity and abundance resulting from *AFG3L2* inactivation closely resemble findings from Lon protease deficient flies (12). In particular, Lon inactivation reduced the abundance of only those RC complexes that contain subunits encoded by the mitochondrial genome. Further studies of Lon-deficient animals revealed that these abundance alterations were caused by a defect in mitochondrial translation (12). To test whether an identical defect in translation explains the RC abundance alterations in Afg3l2-deficient animals, we used 35S-labeled methionine to conduct an *in organello* labeling experiment. This experiment revealed that mitochondrial translation products were nearly absent in pupae expressing the strong *AFG3L2-R2* construct under the control of a da-Gal4 driver (Figure 5A). To explore the mechanism underlying this defect, we subjected mitochondrial protein homogenates from Afg3l2-deficient animals and controls to sucrose density gradient analysis to compare polysome profiles in these genotypes. Use of an antiserum to Mrps22, a component of the small mitochondrial ribosomal subunit, revealed that Mrps22 was only present in fractions 6-8 of the sucrose gradient in Afg3l2-deficient pupae, corresponding to the sedimentation of the 28S small mitochondrial ribosomal subunit (Figure 5B and Supplemental Figure 10). By contrast, Mrps22 was also detected in fractions 12-15 of the sucrose gradient in controls, diagnostic of fully assembled translating 55S monosomes and polysomes (Figure 5B and Supplemental Figure 10). Together, these findings indicate that mitochondrial translation is impaired in Afg3l2-deficient pupae.

**Figure 5.**
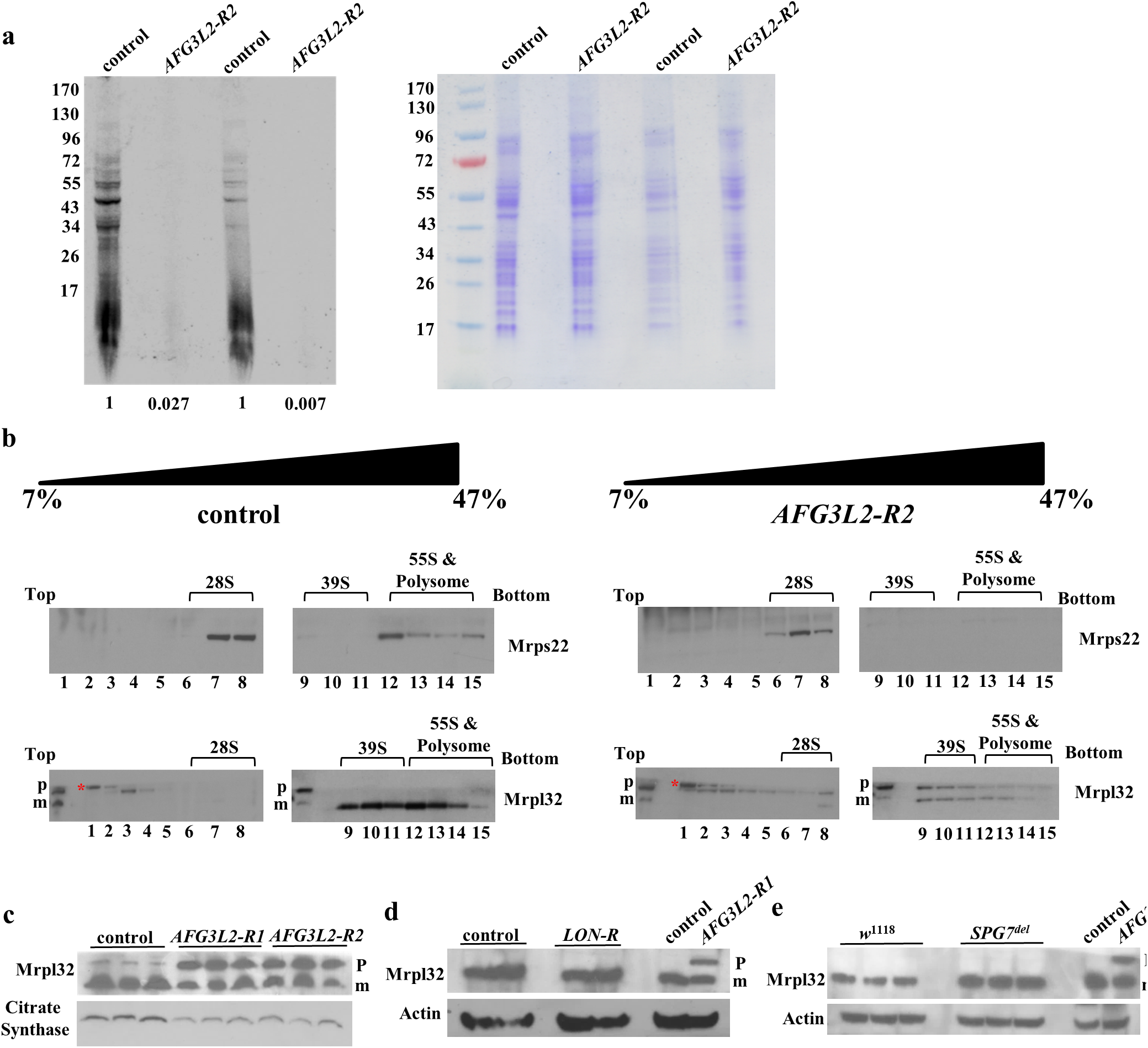
Afg3l2 deficiency impairs mitochondrial ribosome assembly and translation. **a.** Mitochondria from control (UAS-*LUCIFERASE* RNAi/da-Gal4) and Afg3l2-deficient (UAS-*AFG3L2-R2* RNAi/da-Gal4) pupae were labeled with ^35^S-methionine for 1 h, then homogenized and subjected to SDS-PAGE. The left panel shows the ^35^S-labeled proteins, while the right panel shows the Coomassie staining pattern using the same samples. The Coomassie-stained samples were used to normalize protein loading (control samples defined as 1.0). **b.** Sucrose density gradient centrifugation was performed using mitochondrial protein extract from control (UAS-*LUCIFERASE* RNAi/da-Gal4) and Afg3l2-deficient (UAS-*AFG3L2-R2* RNAi/da-Gal4) animals and fractions were subjected to immunoblotting using antisera against Mrps22 and Mrpl32. The red asterisk denotes the non-specific band. **c.** Mrpl32 processing was analyzed by subjecting mitochondrial protein extract from control (UAS-*LUCIFERASE* RNAi/da-Gal4) and Afg3l2-deficient (UAS-*AFG3L2-R1* RNAi; da-Gal4 or UAS-*AFG3L2-R2* RNAi/da-Gal4) pupae to western blot analysis using an antiserum against Mrpl32. Citrate synthase was used as a loading control. The **‘p’** and **‘m’** denote the pre-sequence containing form of Mrpl32 and the mature form of Mrpl32 lacking the presequence, respectively. **d.** Mrpl32 processing was analyzed as in (c) using protein extracts from control (UAS-*LUCIFERASE* RNAi/da-Gal4) and Lon-deficient (UAS-*LON-R* RNAi/da-Gal4) pupae. **e.** Mrpl32 processing was assessed as in (c) and (d) using protein extracts from control (*w*^1118^) and *SPG7^del^* flies. The cell lysate from Afg3l2-deficient (UAS-*AFG3L2-R1* RNAi; da-Gal4) pupae was used as a positive control in **(d)** and **(e)** which exhibited both ‘p’ the pre-sequence containing form of Mrpl32 and ‘m’ the mature form of Mrpl32 lacking the presequence. Actin was used as a loading control in **(d)** and **(e)**.

While our results indicate that translation is impaired in Afg3l2-deficient animals, the nearly complete absence of labeled products from our *in organello* labeling experiment stands in stark contrast to the mild translational defect of Lon-deficient animals, suggested that additional factors may be contributing to the gene expression defect of Afg3l2-deficient animals (12). One possible explanation for the more severe phenotype of Afg3l2-deficient animals derives from previous work in yeast showing that the maturation of Mrpl32, a component of the large subunit of the mitochondrial ribosome, requires a proteolytic processing event that is catalyzed by the *m-*AAA protease orthologs, Yta10 and Yta12 (30). Yeast lacking these *m-*AAA proteases is unable to perform this processing event and exhibit a profound mitochondrial translation defect. To test whether a similar defect in Mrpl32 maturation contributes to the severe translational impairment of Afg3l2-deficient animals, we generated an antiserum to the *Drosophila* Mrpl32 protein and used it to explore the proteolytic processing of Mrpl32 and its assembly into actively translating ribosomes. We found that the efficiency of conversion of full-length Mrpl32 into a shorter processed form was greatly reduced in Afg3l2-deficient pupae relative to controls (Figure 5C). This processing defect was not observed in pupae expressing an RNAi against Lon (*LON-R*) (Figure 5D), or in *SPG7^del^* mutants (Figure 5E) indicating that it is specific to Afg3l2-deficient animals. We also found that the unprocessed and fully processed forms of Mrpl32 were differentially distributed in a sucrose density gradient. The fully processed form of Mrpl32 was detected in two portions of the gradient, diagnostic of the large 39S ribosomal subunit (fractions 9-11) and actively translating monosomes and polysomes (fractions 12-15). By contrast, the unprocessed form of Mrpl32 was found in gradient fractions representing translationally inactive ribonucleoparticles (fractions 1-4), as well as in several higher molecular weight fractions (fraction 6-11) of the gradient (Figure 5B and Supplemental Figure 10). Our finding that the unprocessed form of Mrpl32 is absent from actively translating ribosomes is consistent with findings from yeast, where the unprocessed form of yeast Mrpl32 was also absent from actively translating ribosomes (30). The physical nature of unprocessed Mrpl32 in the higher molecular weight fractions is unclear but may consist of aggregated forms of Mrpl32 that retain the unprocessed pre-sequence. Our findings establish that Afg3l2 protease is required for the processing of Mrpl32 in *Drosophila*, and that a defect in this process likely contributes to the translation defect of Afg3l2-deficient animals.

To test whether impaired translation fully accounts for the reduction in mitochondrially encoded translation products, we also examined mitochondrial DNA (mtDNA) and mitochondrial RNA content in Afg3l2-deficient animals using qPCR. Our analyses did not detect a difference in mtDNA content between Afg3l2-deficient animals and controls (Figure 6A) but did reveal substantial decreases in the abundance of all mtDNA encoded transcripts in Afg3l2-deficient animals, including mRNAs, rRNAs, and tRNA (Figure 6B). To determine if the decreased abundance of mitochondrial transcripts is caused by a defect in transcription or a defect in the post-transcriptional stability of mitochondrial transcripts, we performed an *in organello* transcription assay (31). This experiment revealed a mild decrease in the rate of mitochondrial transcription in Afg3l2-deficient animals (Figure 6C). However, this mild decrease in mitochondrial transcription appears insufficient to fully account for the severe reduction in the abundance of mtDNA encoded transcripts in Afg3l2-deficient animals, suggesting that the reduction in mitochondrial transcript abundance in Afg3l2-deficient animals derives from both reduced mitochondrial transcription and reduced stability of mtDNA encoded transcripts.

**Figure 6.**
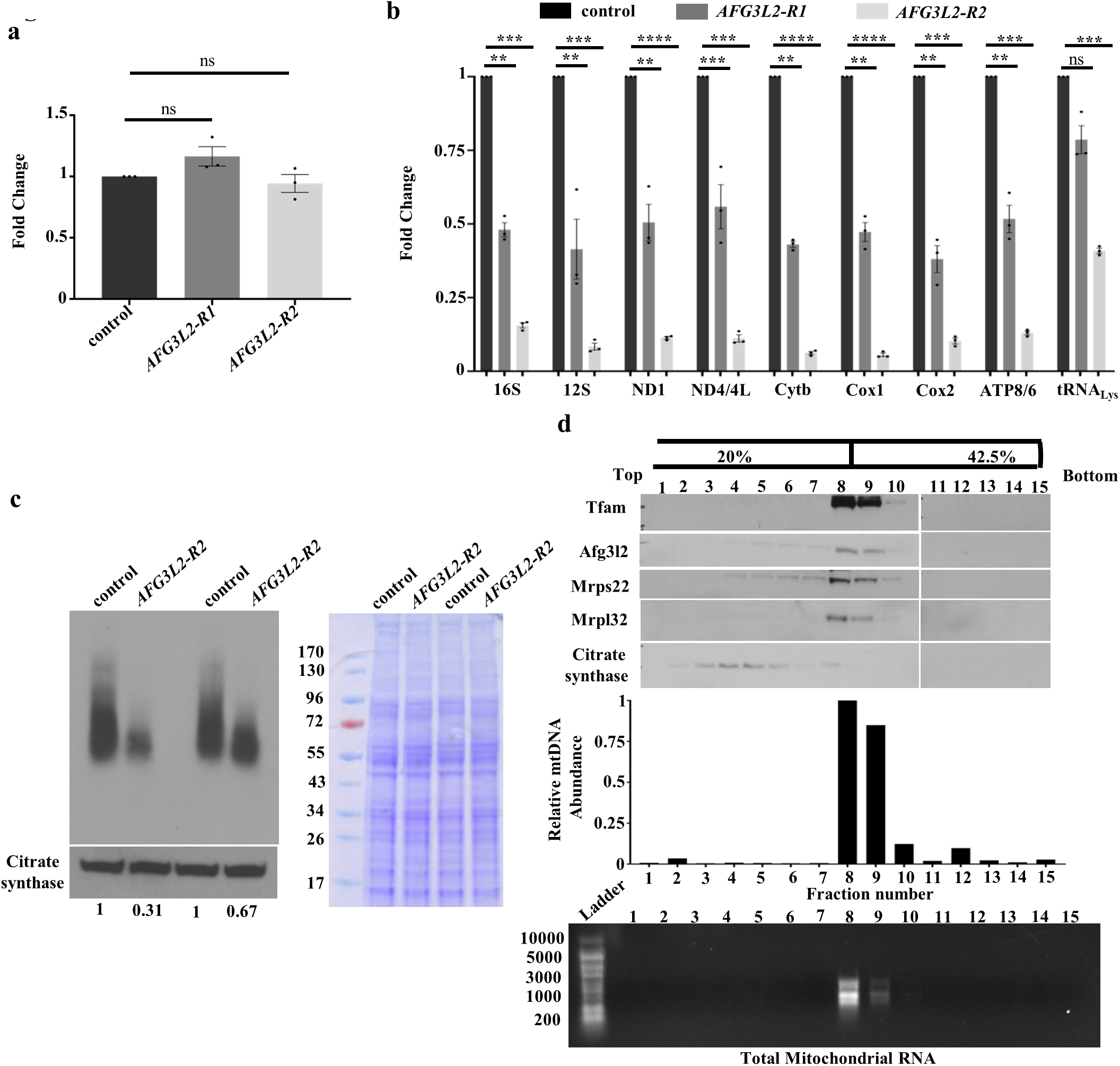
Afg3l2 deficiency impairs mitochondrial transcription. **a.** mtDNA abundance was compared in control (UAS-*LUCIFERASE* RNAi/da-Gal4) and Afg3l2-deficient (UAS-*AFG3L2-R1* RNAi; da-Gal4 and UAS-*AFG3L2-R2* RNAi/da-Gal4) pupae using RT-PCR (N = 3 independent groups of 5 pupae). Statistical significance was determined by one way ANOVA Tukey’s test for multiple comparison. **b.** Quantification of mitochondrial RNA levels in control (UAS-*LUCIFERASE* RNAi/da-Gal4) and Afg3l2-deficient (UAS-*AFG3L2-R1* RNAi; da-Gal4 and UAS-*AFG3L2-R2* RNAi/da-Gal4) pupae using qRT-PCR (N = 3 independent groups of 5 pupae, \*\**p* < 0.005, \*\*\**p* < 0.0005, \*\*\*\**p* < 0.0001 by one way ANOVA Tukey’s test for multiple comparison). **c.** Right panel shows results of *in organello* transcription analysis using mitochondria from control (UAS-*LUCIFERASE* RNAi/da-Gal4) and Afg3l2-deficient (UAS-*AFG3L2-R2* RNAi/da-Gal4) pupae. An antiserum to citrate synthase was used for normalization. Densitometric values after normalization to citrate synthase are indicated below each lane. Right panel shows a Coomassie-stained gel from the same mitochondrial preparation used for *in organello* transcription analysis. **d.** Mitochondrial protein extract isolated from control (UAS-*LUCIFERASE* RNAi/da-Gal4) pupae were subjected to iodixanol density gradient analysis. Fractions from the gradient were subjected to western blot analysis (top panel) using the indicated antisera, qPCR to quantify mtDNA abundance (middle panel), or denaturing agarose gel electrophoresis (bottom panel) to identify mitochondrial RNA-containing fractions.

Our finding that Afg3l2-deficient animals exhibit both a reduction in mitochondrial transcript abundance along with a defect in mitochondrial ribosome biogenesis closely resemble recent work on Letm1, which associates with the mitochondrial nucleoprotein complex and is implicated in ribosome biogenesis (32). To test whether Afg3l2 also associates with the mitochondrial nucleoprotein complex, we subjected detergent-solubilized mitochondrial lysates to iodixanol gradient analysis and examined the distribution of Afg3l2 within the gradient. This analysis revealed that Afg3l2 co-fractionated with mitochondrial DNA and RNA on the iodixanol gradient along with the other components of the mitochondrial nucleoprotein complex, including Tfam, Mrps22, and Mrpl32 (Figure 6D and Supplemental Figure 11). A major fraction of the unrelated protein citrate synthase was present in the early fractions and did not cofractionate with the mitochondrial DNA and RNA (Figure 6D and Supplemental Figure 11). Our findings, therefore, suggest that Afg3l2 is a constituent of the mitochondrial nucleoprotein complex and participates in the mitochondrial RNA metabolism and the assembly of mitochondrial ribosomes.

### Afg3l2 deficiency triggers mitochondrial protein aggregation and activation of mitochondrial stress pathways

The mitochondrial AAA^+^ protease family is thought to play an important role in degrading damaged and misfolded mitochondrial proteins. To test whether Afg3l2 also functions in this capacity, we compared the abundance of mitochondrial protein aggregates in Afg3l2-deficient animals and controls using a previously described detergent extraction method (33, 34). We found that the abundance of many mitochondrial proteins was elevated in detergent-insoluble protein fractions from Afg3l2-deficient pupae, including respiratory chain complex proteins and transcription factors (Figure 7A). We also found that mitochondrial stress pathways that respond to unfolded mitochondrial proteins and mitochondrial dysfunction were activated in Afg3l2-deficient pupae (35). In particular, the mito-UPR markers Lon, Hsp60, and Hsc70-5 were significantly increased in abundance in Afg3l2-deficient pupae (Figure 7B). We also detected induction of markers associated with autophagy and mitophagy in Afg3l2-deficient pupae, including the *Drosophila* LC3 and P62 orthologs, ATG8a and Ref(2)p, respectively (Figure 8A and B) (36–38). This was further accompanied by an increase in total ubiquitinated protein levels (Figure 8B). In addition, we observed an accumulation of Ref(2)p puncta in the Afg3l2-deficient pupae (Figure 8C). These findings indicate that mitochondrial protein unfolding and ensuing mitochondrial dysfunction in Afg3l2-deficient animals triggers activation of compensatory stress pathways designed to alleviate this stress.

**Figure 7.**
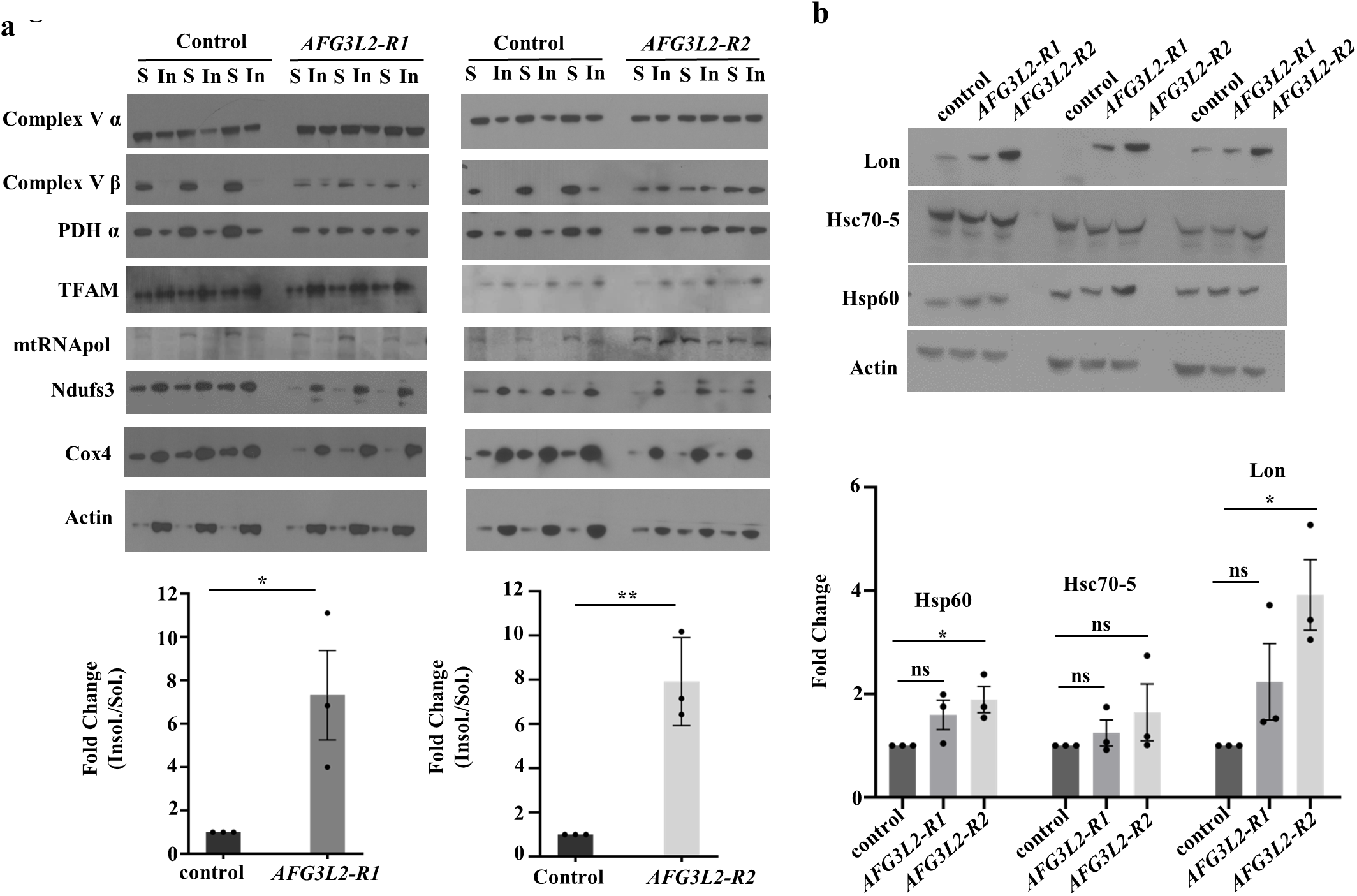
Afg3l2 deficiency results in the accumulation of insoluble mitochondrial proteins and activation of the mito-UPR. **a.** Results of western blot analysis of triton-soluble (S) and insoluble (In) mitochondrial protein fractions from control (UAS-*LUCIFERASE* RNAi/da-Gal4) and Afg3l2-deficient (*AFG3L2-R1*; UAS-*AFG3L2-R1* RNAi; da-Gal4 and UAS-*AFG3L2-R2* RNAi/da-Gal4) pupae using the indicated antisera. The average of all insoluble to soluble proteins was quantified by normalizing band intensities to actin (N = 3 independent biological replicates, \**p* < 0.05, \*\**p* < 0.005 by Student’s t-test). **b**. Results of western blot analysis of cell lysates from control (UAS-*LUCIFERASE* RNAi/da-Gal4) and Afg3l2-deficient (UAS-*AFG3L2-R1* RNAi; da-Gal4 and UAS-*AFG3L2-R2* RNAi/da-Gal4) pupae using antisera to the mito-UPR markers Lon, Hsp60A and Hsc70-5. Quantification was performed after normalizing band intensity of each protein to actin (N = 3 independent groups of 5 pupae, \**p* < 0.05 by one-way ANOVA Tukey’s test for multiple comparison).

**Figure 8.**
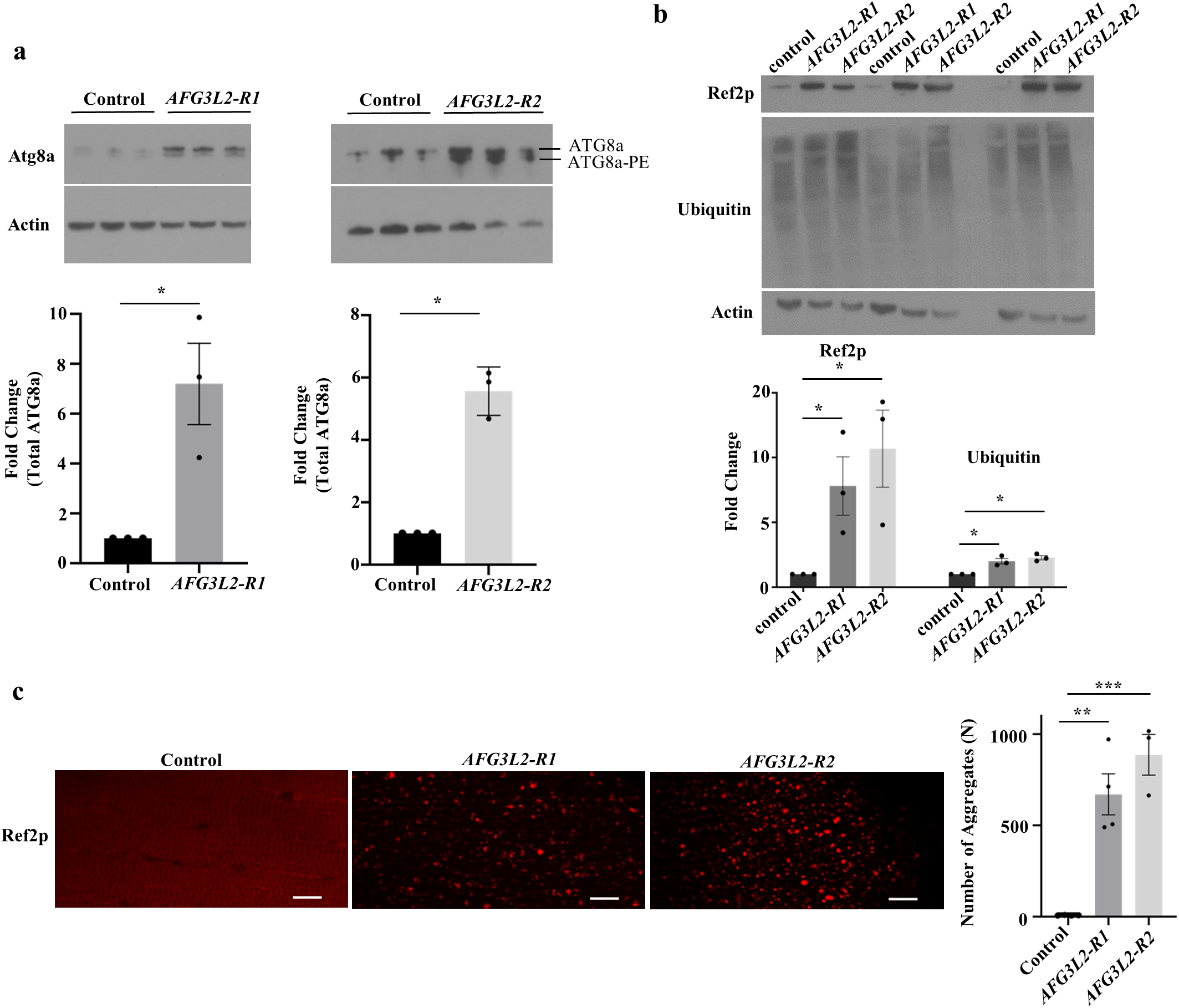
Macroautophagy is induced in Afg3l2-deficient animals. Western blot analysis of cell lysates from control (UAS-*LUCIFERASE* RNAi/da-Gal4) and Afg3l2-deficient (UAS-*AFG3L2-R1* RNAi; da-Gal4 and UAS-*AFG3L2-R2* RNAi/da-Gal4) pupae using antiserum to ATG8a **(a)**, Ref(2)p **(b)** and ubiquitin **(b).** The lower of the two bands detected by the anti-ATG8a antiserum (designated ATG8a-PE) represents a phosphatidylethanolamine conjugated form of Atg8a, whereas the upper band represents the unconjugated form of the ATG8a. Antiserum against Actin was used as a loading control (N = 3 independent groups of 5 pupae, \**p* < 0.05 by one-way ANOVA Tukey’s test for multiple comparison). **c.** Confocal images of indirect flight muscles from Control (UAS-*LUCIFERASE* RNAi/da-Gal4) and Afg3l2-deficient (UAS-*AFG3L2-R1* RNAi; da-Gal4 and UAS-*AFG3L2-R2* RNAi/da-Gal4) pupae using antisera against Ref(2)p. Scale bars are 5 μm. The right panel indicates the average number of Ref(2)p punctate (N = 4 for control and *AFG3L2-R1* animals and N = 3 for *AFG3L2-R2* animals, \*\**p* < 0.005 and \*\*\**p* < 0.0005 by one-way ANOVA Tukey’s test for multiple comparison).

### Activation of the mito-UPR pathway partially rescues the Afg3l2-deficient phenotypes

Our finding that mitochondrial proteins, including mitochondrial transcription factors, exhibited increased aggregation in Afg3l2-deficient animals raised the possibility that sequestration of these factors into inactive aggregates may partially account for phenotypes documented in our work. To test this hypothesis, we examined whether genetic manipulations of mitochondrial chaperones and proteases would influence the phenotypes of Afg3l2-deficient animals. First, we tested whether overexpression of the chaperones Hsp60 or Hsc70-5 or the AAA^+^ protease Lon would suppress the Afg3l2-deficient phenotypes. We found that overexpression of Hsp60, Hsc70-5 and Lon, all partially ameliorated the locomotor defect of Afg3l2-deficient animals (Figure 9A-C). Additionally, overexpression of Hsc70-5 and Lon partially rescued the lifespan defect of Afg3l2-deficient animals (Figure 9D and Supplemental Table 1). These experiments indicate that the phenotypes caused by Afg3l2 deficiency in *Drosophila* are in part reversible by measures that combat protein aggregation. Next, we tested whether inactivating Lon protease would enhance an Afg3l2-deficient phenotype. To perform this experiment, we used the ey-Gal4 driver to express the *AFG3L2-R1* transgene and an RNAi construct targeting Lon (*LON-R*) in the fly eye. Knockdown of either *AFG3L2* or *LON* alone had little or no influence on eye morphology relative to controls (Figure 9E). By contrast, flies coexpressing the *AFG3L2-R1* and *LON-R* transgenes exhibited a substantial defect in eye morphology (Figure 9E). Altogether, our report with Afg3l2-deficient animals demonstrates that accumulation of unfolded proteins account for the phenotypes of Afg3l2-deficient animals.

**Figure 9.**
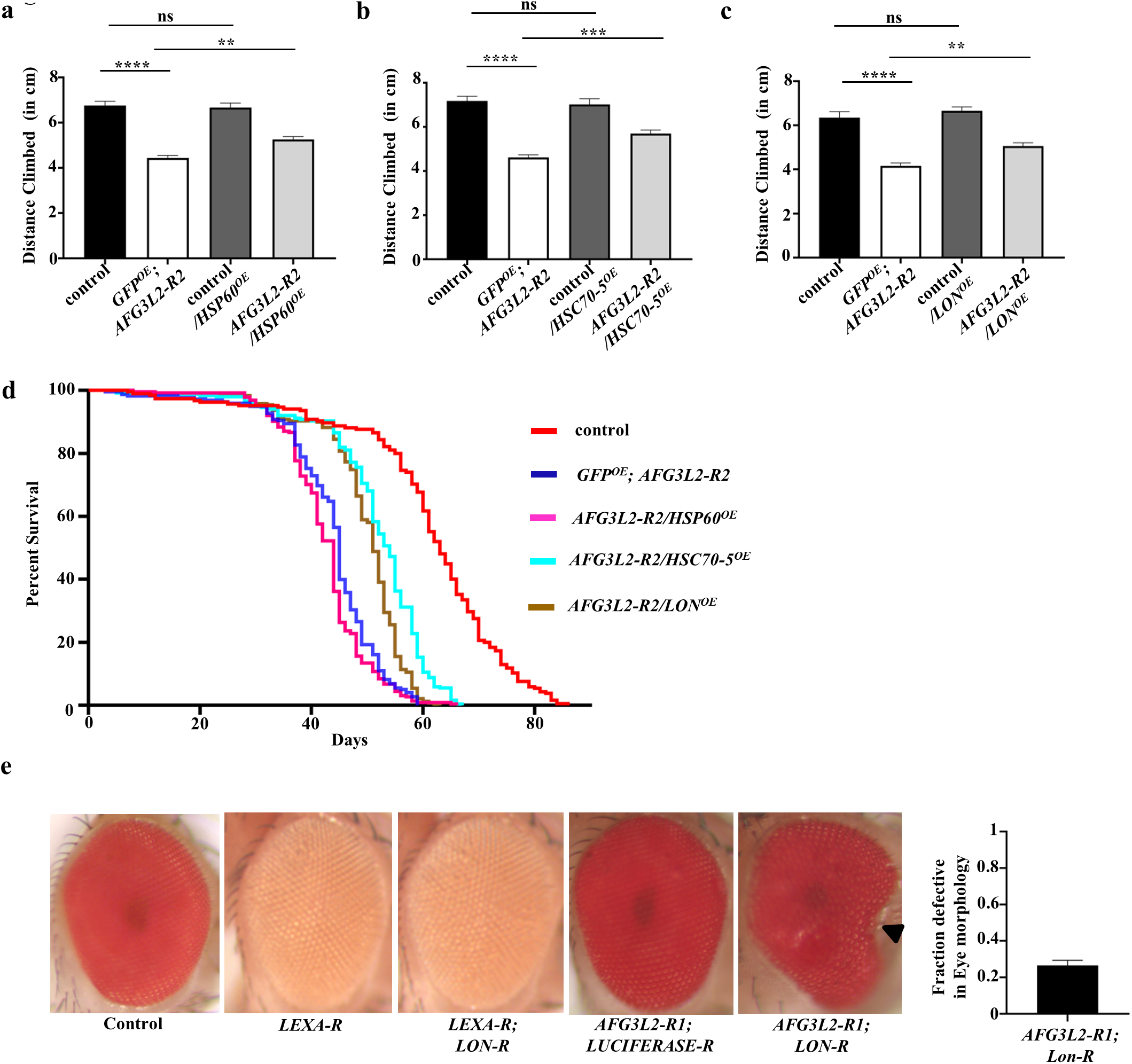
Overexpression of mito-UPR components partially rescues the locomotion and lifespan defects of Afg3l2-deficient animals. **a.** Climbing performance of 1-day old adult controls (elav-Gal4/UAS-*GFP*; UAS-*LUCIFERASE* RNAi; N = 101), Afg3l2-deficient animals: *GFP^OE^; AFG3L2-R2* (elav-Gal4/UAS-*GFP*; UAS-*AFG3L2-R2* RNAi; N = 94), control flies overexpressing Hsp60: control*/HSP60^OE^* (elav-Gal4; UAS-*LUCIFERASE* RNAi*/*UAS-*HSP60*; N = 92) and Afg3l2-deficient animals overexpressing Hsp60: *AFG3L2-R2*/HSP60^OE^ (elav-Gal4; UAS-*AFG3L2-R2* RNAi*/*UAS-*HSP60*; N = 100) (\*\**p* < 0.005, \*\*\*\**p* < 0.0001 by one way ANOVA Tukey’s test for multiple comparison). **b.** Climbing performance of 1-day old adult controls (elav-Gal4/UAS-*GFP*; UAS-*LUCIFERASE* RNAi; N = 63), Afg3l2-deficient animals: *GFP^OE^*; *AFG3L2-R2* (elav-Gal4/UAS-*GFP*; UAS-*AFG3L2-R2* RNAi; N = 65), control flies overexpressing Hsc70-5: Control*/HSC70-5^OE^* (elav-Gal4; UAS-*LUCIFERASE* RNAi*/*UAS-*HSC70-5*; N = 48) and Afg3l2-deficient animals overexpressing Hsc70-5: *AFG3L2-R2*/*HSC70-5^OE^* (elav-Gal4; UAS-*AFG3L2-R2* RNAi*/*UAS-*HSC70-5*; N = 66) (\*\*\**p* < 0.0005, \*\*\*\**p* < 0.0001 by one way ANOVA Tukey’s test for multiple comparison). **c.** Climbing performance of 1-day old adult controls (elav-Gal4/UAS-*GFP*; UAS-*LUCIFERASE* RNAi; N = 53),), Afg3l2-deficient animals: *GFP^OE^; AFG3L2-R2* (elav-Gal4/UAS-*GFP*; UAS-*AFG3L2-R2* RNAi; N = 64), control flies overexpressing Lon: control/*LON*^OE^ (elav-Gal4/*TRiP-OE LON*; UAS-*LUCIFERASE* RNAi/UAS-*dCAS9-FLAG-VPR*; N = 74) and Afg3l2-deficient animals overexpressing Lon: *AFG3L2-R2*/*LON^OE^* (elav-Gal4/*TRiP-OE LON*; UAS-*AFG3L2-R2* RNAi /UAS-*dCAS9-FLAG-VPR*; N = 86). (\*\**p* < 0.005, \*\*\*\**p* < 0.0001 by one way ANOVA Tukey’s test for multiple comparison). **d.** Kaplan–Meier survival curves of controls (elav-Gal4/UAS-*GFP*; UAS-*LUCIFERASE* RNAi; N = 185, 50% survival 67 days), Afg3l2-deficient animals: *GFP^OE^*; *AFG3L2-R2* (elav-Gal4/UAS-*GFP*; UAS-*AFG3L2-R2* RNAi; N = 218, 50% survival 45 days), Afg3l2-deficient animals overexpressing Hsp60: *AFG3L2-R2/HSP60*^OE^ (elav-Gal4; UAS-*AFG3L2-R2* RNAi*/*UAS-*HSP60*; N = 224, 50% survival 44 days), Afg3l2-deficient animals overexpressing Hsc70-5: *AFG3L2-R2*/*HSC70-5^OE^* (elav-Gal4; UAS-*AFG3L2-R2* RNAi*/*UAS-*HSC70-5*; N = 237, 50% survival 52 days) and Afg3l2-deficient animals overexpressing Lon: *AFG3L2-R2/LON^OE^* (elav-Gal4/*TRiP-OE LON*; UAS-*AFG3L2-R2* RNAi/UAS-*dCAS9-FLAG-VPR*; N = 239, 50% survival 51 days). Significance was determined using a Mantel-Cox log-rank test (\*\*\*\**p* < 0.0001). For the Figs. **a-d,** to account for the possible titration of Gal4 protein in the presence of two UAS transgenes and subsequent reduction in expression of the RNAi targeting *AFG3L2*, we included the UAS-GFP transgene along with the UAS-*AFG3L2-R2* RNAi transgene in control animals. **e.** Eye morphology of controls (ey-Gal4; UAS-*LUCIFERASE* RNAi; N = 138), *LEXA-*RNAi expressing flies (ey-Gal4/UAS-*LEXA* RNAi; N = 108), Lon-deficient flies: *LEXA-R*; *LON-R* (ey-Gal4/UAS-*LEXA* RNAi; UAS-*LON* RNAi; N = 169), Afg3l2-deficient animals: *AFG3L2-R1; LUCIFERASE-R* (ey-Gal4/UAS-*AFG3L2-R1* RNAi; UAS-*LUCIFERASE* RNAi; N = 231) and Afg3l2-deficient animals expressing an RNAi construct targeting Lon: *AFG3L2-R1*; *LON-R* (ey-Gal4/UAS-*AFG3L2-R1* RNAi; UAS-*LON* RNAi; N = 241). To control for the non-specific interactions, the control animals included RNAi lines targeting LexA and Luciferase, neither of which are encoded in the *Drosophila* genome. The right panel indicates the quantification of defective eye morphology in *AFG3L2-R1*; *LON-R* flies and none of the other genotypes tested had eye defects.

## DISCUSSION

Mitochondria play a variety of essential cellular functions, but their unique features make them particularly prone to damage. In particular, the fact that the RC complexes are comprised of subunits encoded by the nuclear and mitochondrial genome can lead to stoichiometric imbalances and protein aggregation (39). Moreover, mitochondria are the primary cellular source of damaging ROS, and consequently, the primary recipients of this damage (2). Therefore it is unsurprising that mutation in genes that function in mitochondrial quality control often cause severe incurable disease syndromes (5, 8). To create a foundation for the development of treatments for these syndromes, we have been using a genetic approach in *Drosophila* to study the mechanisms underlying defects in some of these genes (10, 11). In our current work, we inactivated the *Drosophila AFG3L2* gene to create models of two of these syndromes: spastic ataxia neuropathy syndrome and spinocerebellar ataxia (15, 40). We found that a null mutation in *AFG3L2* results in early larval lethality. However, by using RNAi to partially inactivate *AFG3L2*, we were able to substantially extend the lethal phase such that we could better characterize the effects of diminished Afg3l2 activity. We found that Afg3l2 deficiency resulted in a shortened lifespan, neurodegeneration, and locomotor deficits. These phenotypes were accompanied by diminished mitochondrial RC activity and bioenergetic failure that appear to be a consequence of an accumulation of unfolded mitochondrial proteins and widespread defects in mitochondrial gene expression. Our findings thus indicate that we have created valid models of spastic ataxia neuropathy syndrome and spinocerebellar ataxia, and our work on these disease models provides novel insight into the pathogenesis of these disorders.

Perhaps the most striking observation from our work on Afg3l2-deficient animals concerns the catastrophic defect in mitochondrial gene expression. While our finding that Afg3l2-deficient animals exhibit defective processing of Mrpl32 was not unexpected given past work in yeast, our study indicates that the gene expression defects also include reduced mitochondrial transcription, transcript stability, and mitochondrial ribosome assembly. These observations were both unexpected and novel. One likely explanation for these defects is that proteins required for transcription, translation, and ribosome assembly are partially sequestered in misfolded detergent-insoluble aggregates in Afg3l2-deficient animals. In support of this model, we found that the key mitochondrial transcription factors, Tfam and mtRNApol were more abundant in the triton insoluble fraction relative to controls. We also found that the phenotypes of Afg3l2-deficient flies were partially rescued by the overexpression of mito-UPR components, providing further evidence that protein aggregation is at least partly responsible for these gene expression defects. However, another possible explanation for the ribosome assembly defect is provided from our finding that Afg3l2 co-sediments with the mitochondrial nucleoprotein complex along with Tfam and subunits of mitochondrial ribosomes. Mitochondrial ribosome assembly is believed to occur at the nucleoprotein complex, thus our finding raises the possibility that Afg3l2 plays a role in this process (41, 42). However, future studies will be required to validate this role of Afg3l2 and to explore the mechanism by which it contributes to ribosome assembly.

Previous work indicates that the vertebrate AAA-protease subunit Paraplegin does not assemble as a homo-oligomer (17). Rather, Paraplegin forms a functional AAA-protease by assembling with Afg3l2 (43). While our data support the existence of this hetero-oligomeric assembly in *Drosophila*, we found that introducing an Afg3l2 null mutation into a Paraplegin null background failed to enhance the phenotype of Paraplegin mutants. While there are multiple interpretations of this finding, one possible interpretation is that the Afg3l2 homomultimers and the Afg3l2/Paraplegin heteromultimers have completely independent substrates and functions. Further evidence in support of this model is provided by our finding that Paraplegin null mutants do not influence Mrpl32 processing, suggesting that this function is exclusively performed by the Afg3l2 homo-oligomer. Moreover, Paraplegin null mutants did not exhibit defects in the activity or abundance of several RC complexes that contain subunits encoded by the mitochondrial genome, indicating that these mutants do not have a general defect in mitochondrial ribosome assembly (11). In contrast to our expectations, the phenotypic consequences of Afg3l2 deficiency closely resemble those exhibited by Lon-deficient animals, suggesting that the functional role of Afg3l2 may more closely resemble that of Lon, than of the Afg3l2/Paraplegin heteromultimers (12).

In summary, our work provides novel insight into the mechanisms underlying the diseases caused by mutations in *AFG3L2*, and provides a solid foundation to further explore these mechanisms and to test hypotheses that derive from this work using the powerful genetic tools available in *Drosophila*. For example, our finding that both autophagy and the mitochondrial unfolded stress pathways are induced in Afg3l2-deficient animals, likely as compensation for loss of a mitochondrial quality control pathway, suggests that activation of these pathways would be beneficial. Moreover, the broad mitochondrial gene expression defects of Afg3l2-deficient animals suggest that methods that activate gene expression, for example, knockdown of negative regulators of mitochondrial transcription or translation, would also be therapeutic. Our current work provides a starting point for testing these models, as well as addressing other matters of interest, including the use of proteomic methods to identify substrates of Afg3l2. Such work should facilitate a better understanding of the biological roles of Afg3l2 and the molecular basis of the diseases caused by mutations in this gene.

## Materials and methods

### Fly stocks and maintenance

All Drosophila stocks and crosses were maintained on standard cornmeal molasses food at 25 °C using a 12-h light/dark cycle. The *w*^1118^, elav-Gal4 (X), elav-Gal4 (II), 24B-Gal4, da-Gal4, Ey-Gal4, UAS-*LEXA* RNAi, UAS-*LON* RNAi, and UAS-*MCU* RNAi lines were obtained from the Bloomington Stock Center (Bloomington, IN, USA). The UAS-*AFG3L2* RNAi constructs, P{KK101663}VIE-260B (designated here as *AFG3L2-R1*) and P{GD3606}v8515 (designated here as *AFG3L2-R2*) were obtained from the Vienna Drosophila Resource Center. The UAS-*LUCIFERASE* RNAi transgenic line was a kind gift from the Norbert Perrimon lab (Harvard Medical School). The UAS-*HSP60* and UAS-*HSC70-5* transgenic lines were purchased from Fly facility, National Center For Biological Sciences, Bengaluru, India. The *LON* gene was overexpressed using guide RNAs generated by the Transgenic RNAi Project for overexpression of genes (TRiP-OE) and dCas9-VP64 activator (dCas9-VPR) (44). The crossing schemes to perform this analysis are depicted in Supplemental Figure 12.

The *AFG3L2* knockout allele was created by CRISPR/Cas9-mediated gene editing according to a published procedure (14). Briefly, we replaced the *AFG3L2* (*CG6512*) coding sequence with DsRed through homology-mediated repair. The following primer sequences were used for guide RNAs targeting the 5’ and 3’ UTR regions of *AFG3L2*: 5’-Guide RNA Sense oligo: 5’-CTTCGCAATCCGGGCGACCGATCGT −3’ Antisense oligo: 5’-AAACACGATCGGTCGCCCGGATTGC −3’ 3’-Guide RNA Sense oligo: 5’- CTTCGAAAACGGGTTTAAACCGTA −3’ Antisense oligo: 5’-AAACTACGGTTTAAACCCGTTTTC −3’

Sequences flanking the *AFG3L2* coding region were amplified from genomic DNA to facilitate homology-directed repair using the following primer sequences: 5’-Homology arm Forward: 5’-CCGGCACCTGCGGCCTCGCGGAAGCAGTACTGTTCTCTACCCAC-3’ Reverse: 5’-CCGGCACCTGCGGCCCTACATCGGTCGCCCGGATTGCGTGGCCACC-3’ 3’-Homology arm Forward: 5’-GGCCGCTCTTCATATGGTTTAAACCCGTTTTCGAAAACAGCAAAAG −3’ Reverse: 5’-CCGGGCTCTTCTGACGCCGGCCGGCAGCCATTTTGCAGGAAAG −3’

Both 5’ and 3’ homology arms were then cloned into the pHD-DsRed-attP vector containing the eye-specific 3xP3 promoter fused with DsRed and the resulting construct was microinjected into Cas9-expressing embryos by a commercial service (Rainbow Transgenic Flies Inc.). Flies bearing the *AFG3L2* deletion were identified by screening the offspring of injected adults for expression of red fluorescence in the eye. Whole-genome sequencing was also performed to verify the correct targeting of the *AFG3L2* gene.

The UAS-*AFG3L2* transgene was PCR amplified using a cDNA clone as a template from *Drosophila* Genomics Resource Center using the following primer sequences: Forward: 5’-GGCCGAATTCACAAAATGGCGTTCCGGCTGCTTGGCACGG −3’ Reverse: 5’-GGCCGCGGCCGCCTAACTGCTCTGGGCAGTTACGGGCTTGG −3’ The Kozak consensus sequence, ACAAA was added to the 5’ for optimum translational efficiency. The PCR product was then cloned into the PUAST vector and the resulting construct was microinjected into *w*^1118^ embryos by a commercial service (Rainbow Transgenic Flies Inc). Flies bearing the mini-white marker were crossed to a da-Gal4 driver line and subjected to immunoblot analysis using an anti-Afg3l2 antiserum to confirm that these lines overexpress Afg3l2 protein. Only fly lines exhibiting overexpression of Afg3l2 were used in subsequent experiments.

### Behavioral analyses

All behavioral analyses were performed using male flies. For Lifespan assays, groups of 10–15 flies were transferred to new vial every second day and the number of dead flies was counted. Kaplan–Meier survival curves were generated using Microsoft Office Excel, and the log-rank test was used to determine statistical significance.

For locomotor analyses, flies were anesthetized by brief exposure to CO2 and, allowed for recovery for 1 day before the experiment. Climbing behavior was assessed using the Rapid Iterative Negative Geotaxis (RING) assay on day 1, according to a previously published protocol (45). Briefly, plastic vials containing 10-15 flies each were loaded onto the RING apparatus. To initiate the climbing response, the apparatus was gently tapped down 3–4 times and the height climbed by each fly after 4 s was measured using ImageJ software.

Age in all experiments refers to the number of days after eclosion for adult flies and days after egg hatching for experiments involving larvae or pupae.

### Histological analysis

Histological analysis of brain tissue was performed as previously described (34). Briefly, brain tissues from 1-day old adult flies were fixed in Carnoy’s fixative (10% acetic acid; 30% chloroform; 60% absolute ethanol) for 3.5 h and dehydrated in ethanol. After paraffin infiltration at 60 °C, 4-μm sections were analyzed by hematoxylin and eosin staining. Images were acquired on a Nikon Optiphot-2 using a 10× objective. Brain vacuole size and area was quantified using ImageJ software.

### Immunoblotting

1-day old adult whole flies/fly heads or 9-day old pupae were homogenized in 1x laemmli buffer and the supernatant was subjected to SDS-PAGE followed by western blotting using primary antisera at the following dilutions: mouse anti-Actin (1:10,000, Chemicon MAB1501), rabbit anti-Afg3l2 (1:1000, Antibody plus CSB-PA04005A0Rb), mouse anti-Ndufs3 (1:1000, abcam ab14711), mouse anti-Cox4 (1:1000, abcam ab33985), mouse anti-ATPVα (1:1000, abcam ab14748), mouse anti-ATPVβ (1:1000, Thermo Fisher Scientific A-21351), rabbit anti-Lon (1:1000, Novus Biologicals NBP1-81734), mouse anti-PDH (1:3000, MitoSciences MSP07), rabbit anti-Tfam (1:1000, Abcam ab47548), rabbit anti-mtRNApol (1:500, aviva systems biology ARP48659_P050), rabbit anti-citrate synthase (1:1000, Alpha Diagnostics CISY11-A), rabbit anti-Hsp60 (1:1000, Cell Signaling Technology D307), rabbit anti-Lrpprc (1:1000, a kind gift from the Paul M. Macdonald laboratory), rabbit anti-Grp75 (1:1000, Santa Cruz sc-13967), rabbit anti-Ref(2)p (1:500, abcam ab178440), mouse anti-ubiquitin (1:1000, Enzo Life Sciences International Inc BML-PW8810-0100), rabbit anti-Atg8a (1:5000, Millipore ABC974), rabbit anti-Paraplegin (1:1000, custom antiserum raised against amino acid 790-813, generated by Yenzym Antibodies LLC), rabbit anti-Mrps22 (1:1000, custom antiserum raised against amino acid 311-328, generated by Yenzym Antibodies LLC) and rabbit anti-Mrpl32 (1:1000, custom antiserum raised against amino acid 173-193, generated by Yenzym Antibodies LLC). Chemiluminescence was used for detection and western blot images were quantified using ImageJ software and normalized to Actin. Each experiment was performed at least three times.

The accumulation of mitochondrial unfolded proteins was analyzed according to previously published protocols (12, 33).

### Immunofluorescence and confocal microscopy

To examine the cellular distribution of Afg3l2, salivary glands from 3^rd^ instar larvae were dissected in cold PBS buffer then fixed in 4% paraformaldehyde for 1 h. After washing with PBS (including 0.3% Triton X-100), dissected tissues were then incubated overnight with rabbit anti-Afg3l2 (1:250, Antibody plus CSB-PA04005A0Rb) and mouse anti-cox4 (1:1000, BD Biosciences) antisera, then washed with PBS. Tissues were then incubated overnight with antirabbit Alexa 488 and anti-mouse Alexa 568 secondary antiserum (both at 1:500) and imaged using an Olympus FV-1000 confocal microscope with a 60x lens and a 3x digital zoom. Each stack of 38 images was deconvoluted using Huygens Professional 4.4.0-p8 software (Scientific Volume Imaging) using a signal to noise ratio of 20 for the red and green channels.

The Ref(2)p aggregates were analyzed using indirect flight muscles dissected from pupae 9 days after egg hatching. After fixing in paraformaldehyde, tissues were incubated with anti-Ref(2)p (1:250) antisera. All washes and secondary antibody incubations were performed as described above.

### Transmission electron microscopy

Transmission electron microscopy was performed as previously described (11). Briefly, thoraces from pupae 9 days after egg hatching were dissected in fixative containing 2.5% glutaraldehyde, and 2% paraformaldehyde in 0.1M sodium cacodylate buffer, pH 7.4. Fixed tissues were then treated with 1% OsO4, dehydrated in an ethanol series, and embedded using Epon. Ultra-thin sections of 70 nm were stained with 6% uranyl acetate and a Reynolds lead citrate solution then examined using a JEOL JEM 1400 transmission electron microscope. Mitochondrial cristae density was calculated using Image J software.

### Mitochondrial respiratory chain activity, total ATP content, Blue Native PAGE (BN-PAGE) and in-gel activity assay

Mitochondrial respiratory chain activity, total ATP content, BN-PAGE, and in-gel activity assays were performed according to a published procedure with minor modifications (12). Briefly, 1 ml larvae 4 days after egg hatching/pupae 9 days after egg hatching per biological replicate were homogenized in 10 ml isolation buffer (5 mM HEPES (pH 7.4), 75 mM sucrose, 225 mM mannitol and 0.1 mM EGTA) with 2 % (w/v) fatty acid-free Bovine Serum Albumin (BSA). The lysate was then passed through a 70 μm nylon mesh to remove cellular debris and subjected to centrifugation at 1500 g for 6 min. The supernatant was next subjected to further centrifugation at 8000 g for 6 min. The mitochondrial pellet was washed twice in isolation buffer without BSA, resuspended in the same buffer, flash-frozen in liquid nitrogen and stored at −80 °C. To measure Complex I activity, 100 μg of mitochondria were incubated in the presence of NADH and ubiquinone-1 and the change in absorbance was monitored spectrophotometrically at 340 nm. Background activity was determined using 10 μM rotenone to inhibit complex I and the absorbance change under these conditions was subtracted from the absorbance change in the absence of rotenone to calculate complex I-specific activity. Complex II activity was measured by monitoring the change in absorbance at 600 nm in the presence of 2,6-dichlorophenolindophenol, succinate, and decylubiquinone. Background activity was calculated using 10 mM malonate. Complex III activity was determined by monitoring the reduction of cytochrome c in the presence of decylubiquinol at 550 nm. Background activity was determined using antimycin A. Complex IV activity assay was performed by monitoring the oxidation of reduced cytochrome c at 550 nm in the presence of 1 μg of mitochondria. Background activity was determined using potassium cyanide. Normalization in all experiments was performed by using citrate synthase activity, calculated by monitoring the reduction of 5, 5-dithiobis (2-nitrobenzoic acid) at 412 nm in the presence of acetyl-coenzyme A and oxaloacetate.

Total ATP content was determined by homogenizing five 9-day-old pupae in 100 μL of buffer containing 6M guanidine HCL, 100 mM Tris (pH 7.8), and 4 mM EDTA. Homogenates were then boiled for 5 min and subjected to centrifugation for 3 min at 21,000 g to remove debris. The supernatant was diluted 1:100 and luminescence was monitored using an ATP determination kit (A22066, Molecular Probes). Luminescence readings for each sample were compared to an ATP standard curve and normalized to the total amount of protein determined using the Pierce BCA Protein Assay Kit (23225, Thermo Scientific).

For BN-PAGE, 100 μg of mitochondria were solubilized using a digitonin/protein (w/w) ratio of 8 and subjected to centrifugation at 20,000 g for 10 min at 4 °C. Coomassie G-250 was added to the supernatant and the sample was analyzed by native PAGE. The combined complex I and IV in-gel activity assay was performed by first incubating the gel in a solution containing cytochrome c and 3,3′-diaminobenzidine in phosphate buffer (pH 7.4) for 40 min. After the appearance of brown reaction products, the gel was washed with water and incubated with the complex I substrate NADH and nitrotetrazolium blue chloride in Tris buffer (pH 7.4) for 20 min. The reaction was quenched with 10% acetic acid upon the appearance of a violet color indicative of complex I activity. For the complex II in-gel activity assay, the gel was incubated in a solution consisting of sodium succinate, nitrotetrazolium blue chloride, and phenazine methosulfate in Tris buffer (pH 7.4) for 40 min. The reaction was quenched with 10% acetic acid upon the appearance of a violet color indicative of complex II activity. The complex V in-gel activity assay was carried out by incubating the gel in a solution containing Tris, glycine, magnesium sulfate, adenosine triphosphate, and lead (II) nitrate for 16 h. The reaction was stopped using 50% methanol upon the appearance of silver bands indicative of complex V activity.

### Co-Immunoprecipitation (Co-IP)

500 μg of mitochondria was solubilized on ice for 20 min in IP buffer (50 mM Tris-HCl (pH 7.4), 150 mM NaCl, 10% glycerol, 10 mM MgCl_2_ and 1 mM ATP) containing 1.5% digitonin. After centrifugation at 21000 g for 20 min, the supernatant was diluted with an IP buffer to achieve 0.5% digitonin and incubated overnight with 50 μg of rabbit anti-Afg3l2 primary antiserum. The lysate was then further incubated overnight with 100 μl of equilibrated protein A beads. After washing three times with IP buffer, elution was performed using 1x laemmli buffer followed by electrophoresis and immunoblotting. Rabbit pre-immune serum was used as a negative control. Veriblot secondary antibody (1:1000, Abcam ab131366) was used to avoid the detection of light and heavy chain bands of IgG.

### Calcium uptake assay

Calcium uptake assays were performed as described previously (26). Briefly, 200 μg of freshly isolated mitochondria from 9-day old pupae were incubated in respiration buffer (250 mM sucrose, 10 mM MOPS-Tris (pH 7.4), 5 mM glutamate, 2.5 mM malate, 5 mM Pi, 0.01 mM EGTA and 0.001 mM of Oregon green BAPTA 6F (Thermo Fisher)). This solution was then subjected to 500 μM pulses of CaCl2 after every 1 minute until saturation in calcium uptake was observed. This experiment was performed using two biological replicates and the representative trace is shown in Fig. 4b.

### Mitochondrial membrane potential measurement

Mitochondrial membrane potential was measured according to previously published protocols with minor modifications (46). Briefly, freshly mitochondria isolated from pupae were resuspended in respiration buffer (0.5 mg/ml) containing Tetramethylrhodamine methyl ester perchlorate (0.5 μM) (Thermo Fisher). The excitation spectra were scanned from 520 nm to 580 nm using 590 nm emission wavelengths. Mitochondrial membrane potential was estimated from the 573/546 fluorescence ratio. To check for specificity, the scan was repeated in the presence of Carbonyl cyanide m-chlorophenyl hydrazine (CCCP) (0.5 μM) to depolarize the mitochondria.

### Mitochondrial DNA and RNA quantification reverse transcription-PCR (qRT-PCR)

Mitochondrial and nuclear DNA was isolated from pupae 9 days after egg hatching using the DNeasy Blood & Tissue kit (Qiagen). A total of 50 ng genomic DNA was used as a template to perform qPCR using iTaq Universal SYBR Green Supermix (Bio-Rad). The mtDNA levels were estimated using PCR primers to amplify the *mt:Cox1* gene and normalizing to the levels of nuclear DNA using primers to amplify the nuclear gene *RAP2L*. The relative fold change was determined using the 2^−ΔΔCt^ method (47).

RNA isolation and reverse transcription-PCR (qRT-PCR) was performed according to a previously published procedure (12). Total RNA was prepared from pupae using the Direct-zol RNA MiniPrep kit (Zymo Research). The iScript cDNA Synthesis Kit (Bio-Rad) was used to reverse transcribe RNA to cDNA. qRT-PCR experiments were performed using iTaq Universal SYBR Green Supermix and a Light Cycler 480 (Roche). Each sample was analyzed in triplicate and normalized to Rap2l transcript abundance. The relative fold change was determined by the 2^−ΔΔCt^ method. All primers used for qRT-PCR are described previously (12) and listed in supplemental table 2.

### *In organello* translation

*In organello* labeling of mitochondrial translation products were performed as previously described (48). 500 μg of mitochondria was isolated from pupae 9 days after egg hatching and resuspended in 500 μl of translation buffer (100 mM mannitol, 10 mM sodium succinate, 80 mM potassium chloride, 5 mM magnesium chloride, 1 mM potassium phosphate, 25 mM HEPES (pH 7.4), 60 μg/ml all amino acids except methionine, 5 mM ATP, 0.2 mM GTP, 6 mM creatine phosphate, and 60 μg/ mL creatine kinase) supplemented with 500 μCi/ml of ^35^S-methionine (Perkin–Elmer). Following incubation at 37 °C for 1 h, mitochondria were washed four times using isolation buffer and resuspended in SDS sample buffer. Roughly 50 μg of mitochondria were subjected to SDS-PAGE. Following electrophoresis, the gel was dried and exposed to a phosphor screen. The phosphor screen was scanned using a gel imaging scanner (GE Typhoon FLA 9000). Roughly 50 μg of mitochondria from the same preparation was loaded on the gel and stained with Coomassie for use as a loading control. The experiment was performed using two biological replicates.

### Mitochondrial ribosome profiling

Mitochondrial ribosomal profiling was performed according to a previously published procedure (48). 2 mg of mitochondria isolated from pupae 9 days after egg hatching and incubated on ice in a lysis buffer containing 260 mM sucrose, 100 mM NH4Cl, 10 mM MgCl_2_, 30 mM Tris-HCl (pH 7.5), 50 U/ml Protector RNase Inhibitor, 100 μg/ml chloramphenicol, 1% Triton X-100 supplemented with EDTA-free complete protease inhibitor cocktail (Roche) and PhosSTOP phosphatase inhibitor cocktail (Roche). Insoluble proteins were pelleted by centrifugation at 9,200 g for 45 min at 4 °C and the supernatant was carefully loaded onto a 7–47% linear sucrose gradient prepared in a buffer containing 100 mM NH4Cl, 10 mM MgCl_2_, 30 mM Tris-HCl (pH 7.5) and EDTA-free complete protease inhibitor cocktail (Roche). Samples were then subjected to ultracentrifugation at 39,000 rpm for 8 h at 4 °C. Fractions of 750 μl were collected by carefully pipetting from the top and proteins were precipitated from each fraction using Trichloroacetic acid – Sodium deoxycholate as previously described (49). Protein samples were then subjected to SDS-PAGE and western blot analysis using antisera against Mrpl32 and Mrps22.

### *In organello* transcription

*In organello* transcription assays were performed on freshly isolated mitochondria as previously described with some modifications (48). Briefly, 250 μg of mitochondria were isolated from pupae 9 days after egg hatching and resuspended in 90 μl of transcription incubation buffer (25 mM sucrose, 75 mM sorbitol, 100 mM KCl, 10 mM K2HPO4, 50 μM EDTA, 5 mM MgCl_2_, 1 mM ADP, 10 mM glutamate, 2.5 mM malate, 10 mM Tris-HCl (pH 7.4) and 1 mg/ml BSA) supplemented with 10 μl of DIG RNA labeling mix (Roche 12039672910) and incubated for 1 h at 37°C with occasional gentle tapping. Following incubation, mitochondria were washed twice with 1 ml of washing buffer (10% glycerol, 10 mM Tris-HCl (pH 6.8) and 0.15 mM MgCl_2_) and RNA was isolated using Trizol Reagent (Invitrogen) according to manufacturer instructions. The isolated RNA was immediately subjected to northern blotting using the northern blot kit (ThermoFisher Scientific AM1940). Following the transfer, the membrane was probed with Anti-Digoxigenin-AP antibody (Millipore Sigma) and the signal was detected using CDP-Star reagent (Millipore Sigma).

### Iodixanol gradient fractionation and separation of the nucleoprotein complex

The mitochondrial nucleoprotein complex isolation using the iodixanol gradient was performed according to a published procedure (32). Briefly, mitochondria were isolated from pupae 9 days after egg hatching in a buffer containing 225 mM mannitol, 75 mM sucrose, 10 mM Tris (pH 7.6), and 1 mM EDTA with 0.1 % (w/v) fatty acid-free Bovine Serum Albumin (BSA). Roughly 2 mg of mitochondria were solubilized in a buffer containing 210 mM mannitol, 70 mM sucrose, 20 mM HEPES (pH 7.8), 2 mM EDTA and 0.4% n-Dodecyl-β-D-Maltoside (DDM) detergent on ice. The lysate was then centrifuged at 1000 g for 10 min and the supernatant was loaded onto a 20-42.5% iodixanol gradient prepared in a buffer containing 20 mM HEPES (pH 7.8), 1 mM EDTA, 50 mM NaCl, 2 mM Dithiothreitol and 0.05% DDM. Samples were then subjected to ultracentrifugation at 28,900 rpm for 14 h at 4 °C. Fractions of 750 μl were collected by carefully pipetting from the top and each fraction was divided into three equal portions. One portion was subjected to protein extraction; a second portion was subjected to DNA isolation; the third portion was subjected to RNA isolation. The protein samples were subjected to SDS-PAGE analysis followed by western blot analysis using antisera to Tfam, Afg3l2, citrate synthase, Mrpl32, and Mrps22. The DNA samples were subjected to mitochondrial DNA quantification as described above. The RNA samples were subjected to denaturing glyoxal gel analysis according to the manufacturer’s instructions (Thermo Fisher).

### Eye severity score

Eye severity was scored using a simple binary metric of eye morphology. Flies with normal eye morphology were given a score of ‘0’ and flies with any defect in the ommatidial organization, or eye size was given a score of ‘1’. All scoring was performed by an investigator blinded to genotype.

### Statistics

All data is represented as mean ± s.e.m. Statistical significance tests were performed using GraphPad Prism 7.

## Acknowledgments

We thank Dr. Scott Kennedy (Department of Pathology, University of Washington) for assistance with whole-genome sequencing of the *AFG3L2^del^* mutant; Kim Miller (Microscopy and Imaging Facility, Virginia Merrill Bloedel Hearing Research Center, University of Washington) for technical assistance with confocal imaging and analysis; Dr. Ying-tzang Tien (Department of Pathology, University of Washington) for histological staining of *Drosophila* tissue sections; Dr. Bobbie Schneider (Fred Hutchinson Cancer Research Center) for assistance with electron microscopy; Dr. Yasemin Sancak lab for allowing us to use of her lab space and reagents for experiments involving calcium uptake; Dr. David R. Morris for allowing us to use of his lab space for experiments involving mitochondrial ribosome profiling; Dr. Dana Miller for allowing us to use of her lab space for experiments involving radioactive isotopes; Dr. Evan Eichler and Dr. Stanley Fields for providing access to their lab facilities; Dr. Alexander Whitworth for providing us with the UAS-*MCU-FLAG*, UAS-*EMRE-MYC,* and UAS-*MICU1-HA* fly stocks, Dr. Norbert Perrimon lab for providing us UAS-*LUCIFERASE* RNAi stock, Dr. Paul M. Macdonald for providing an antiserum against the Lrpprc protein and Dr. Ruth E Thomas and Dr. Evelyn Vincow from Pallanck lab for critical review of this work and manuscript. This work was supported by a grant from the National Institute of Health to LP (R21NS094901).

## Supporting Information Legends

### Supporting Figure Legends

**Fig. S1 *Drosophila* Afg3l2 localizes to mitochondria.**

**a.** Confocal image of salivary glands from third instar larvae using antisera against Afg3l2 (left panel) and Cox4 (middle panel) and a merged image (right panel) showing the degree of Afg3l2 colocalization with Cox4. The scale bar is 5 μm. **b.** Western blot of mitochondrial and cytosolic fractions from adult flies using antisera against Afg3l2, Cox4, and Actin.

**Fig. S2 Ectopic expression of *AFG3L2* rescues the lethality of *AFG3L2^del^* homozygotes.**

**a.** Western blot analysis of whole fly extracts control flies and from flies bearing a UAS-*AFG3L2* transgene and the da-Gal4 driver. **b.** Image showing a control fly and a viable adult *AFG3L2^del^* homozygote rescued by ectopic expression of Afg3l2 driven by the da-Gal4 driver (*AFG3L2^del^* da-Gal4 > UAS-*AFG3L2*).

**Fig. S3 RNAi lines targeting *AFG3L2* severely diminish Afg3l2 protein abundance.**

**a.** Western blot analysis of protein lysates prepared from controls (UAS-*LUCIFERASE* RNAi/da-Gal4) and Afg3l2-deficient (UAS-*AFG3L2-R1* RNAi; da-Gal4 and UAS-*AFG3L2-R2* RNAi/da-Gal4) pupae 9 days following egg hatching. **b.** Western blot analysis of a head protein extract from 1-day old adult controls (elav-Gal4; UAS-*LUCIFERASE* RNAi) and age-matched flies expressing RNAi constructs targeting Afg3l2 using the neuron-specific elav-Gal4 driver (elav-Gal4; UAS-*AFG3L2-R1* RNAi and elav-Gal4; UAS-*AFG3L2-R2* RNAi). Band intensities were normalized to actin (N = 3, \*\*\*\**p* < 0.0001 by one-way ANOVA Tukey’s test for multiple comparison).

**Fig. S4 Quantification of the number and area of brain vacuoles in Afg3l2-deficient flies.** The number (**a**) and area (**b**) of brain vacuoles in 1-day old adult fly heads of controls (elav-Gal4; UAS-*LUCIFERASE* RNAi) and flies expressing the weaker RNAi line targeting *AFG3L2* (elav-Gal4; UAS-*AFG3L2-R1* RNAi) throughout the nervous system (N = 6 fly heads, *p < 0.05, ***p <0.0005 by Student’s t-test).

**Fig. S5 Respiratory chain activity is diminished in Afg3l2-deficient larvae.**

RC activity was analyzed using mitochondria isolated from control (UAS-*LUCIFERASE* RNAi/da-Gal4) and Afg3l2-deficient (UAS-*AFG3L2-R1* RNAi; da-Gal4 and UAS-*AFG3L2-R2* RNAi/da-Gal4) 3^rd^ instar larvae (N = 3 independent biological replicates, *p < 0.05, **p < 0.005 by one-way ANOVA Tukey’s test for multiple comparisons).

**Fig. S6 Afg3l2-deficient animals do not accumulate oxidatively damaged proteins.**

Protein extracts from control (UAS-*LUCIFERASE* RNAi/da-Gal4) and Afg3l2-deficient (UAS-*AFG3L2-R1* RNAi; da-Gal4 and UAS-*AFG3L2-R2* RNAi/da-Gal4) pupae were subjected to western blot analysis using an antiserum to Dinitrophenyl to detect protein carbonylation. Band intensities were normalized to actin. Significance was determined using one-way ANOVA Tukey’s test for multiple comparisons.

**Fig. S7 Respiratory chain complex subunits exhibit reduced abundance in Afg3l2-deficient animals.**

Cell lysates from control (UAS-*LUCIFERASE* RNAi/da-Gal4) and Afg3l2-deficient (UAS-*AFG3L2-R1* RNAi; da-Gal4 and UAS-*AFG3L2-R2* RNAi/da-Gal4) pupae were subjected to western blot analysis using the indicated antisera. Band intensities were normalized to actin (N = 3 independent biological replicates, \**p* < 0.05, \*\**p* < 0.005 by one way ANOVA Tukey’s test for multiple comparison).

**Fig. S8 Complex V assembly is defective in Afg3l2-deficient animals.**

Mitochondrial protein extracts from control (UAS-*LUCIFERASE* RNAi/da-Gal4) and Afg3l2-deficient (UAS-*AFG3L2-R1* RNAi; da-Gal4 and UAS-*AFG3L2-R2* RNAi/da-Gal4) pupae were subjected to BN-PAGE analysis. A sub-complex containing the F1 subunit of ATP synthase was detected Afg3l2 animals, but not in controls. Citrate synthase was used as a loading control.

**Fig. S9 Afg3l2-deficiency results in increased MCU abundance.**

**a.** Cell lysates prepared from control (UAS-*LEXA* RNAi; UAS-*EMRE-MYC*/da-Gal4) and Afg3l2-deficient (UAS-*AFG3L2-R1* RNAi; UAS-*EMRE-MYC*/da-Gal4) pupae expressing a Myc-tagged form of EMRE were subjected to western blot analysis using an antiserum against Myc. **b.** Cell lysates prepared from control (UAS-*LEXA* RNAi; UAS-*MICU1-HA*/da-Gal4) and Afg3l2-deficient (UAS-*AFG3L2-R1* RNAi; UAS-*MICU1-HA*/da-Gal4) pupae expressing an HA-tagged form of Micu1 were subjected to western blot analysis using an antiserum against HA. **c.** Cell lysates prepared from control (UAS-*MCU-FLAG*; UAS-*LUCIFERRASE* RNAi/da-Gal4) and Afg3l2-deficient (UAS-*MCU-FLAG*; UAS-*AFG3L2-R2* RNAi/da-Gal4) pupae expressing a Flag-tagged form of Mcu were subjected to western blot analysis using an antiserum against Flag. Band intensities were normalized against actin (N = 3 independent biological replicates, *p < 0.05 by Student’s t-test).

**Fig. S10 Mitochondrial ribosome assembly is impaired in Afg3l2-deficient animals.**

Mitochondrial protein fractions from controls (UAS-*LUCIFERASE* RNAi/da-Gal4) and Afg3l2 deficient (UAS-*AFG3L2-R2* RNAi/da-Gal4) pupae were subjected to sucrose density gradient sedimentation. Individual fractions from the gradient were subjected to western blot analysis using antisera against Mrps22 and Mrpl32. The red asterisk denotes the non-specific band. Data shown represent independent biological replicates.

**Fig. S11 Characterization of an Afg3l2-containing mitochondrial nucleoprotein complex.**

A mitochondrial protein lysate from control (UAS-*LUCIFERASE* RNAi/da-Gal4) pupae was subjected to iodixanol density gradient analysis. Fractions from the gradient were then subjected to western blot analysis using antisera against Tfam, Afg3l2, citrate synthase, Mrpl32, and Mrps22.

**Fig. S12 The crossing schemes used to generate flies overexpressing Hsp60, Hsc70-5, and Lon in control and Afg3l2-deficient backgrounds.**

### Supporting Table Legends

**Supplemental Table 1 Summary of Lifespan analyses for the indicated genotypes.**

Note that two different elav-Gal4 transgenes were utilized in our studies. The elav-Gal4 transgene situated on the ‘X’ chromosome (Elav (X)) was used for all initial characterization of UAS-*AFG3L2-R1* and UAS-*AFG3L2-R2* RNAi flies. The elav-Gal4 transgene on the 2^nd^ chromosome (Elav (II)) was utilized for all overexpression studies.

**Supplemental Table 2 Primers for qRT-PCR analysis**

