## SUPPLEMENTAL FIGURES AND TABLES for "Inactivation of the mitochondrial protease Afg3l2 results in severely diminished respiratory chain activity and widespread defects in mitochondrial gene expression"

### Supplemental Figure 1

**a**

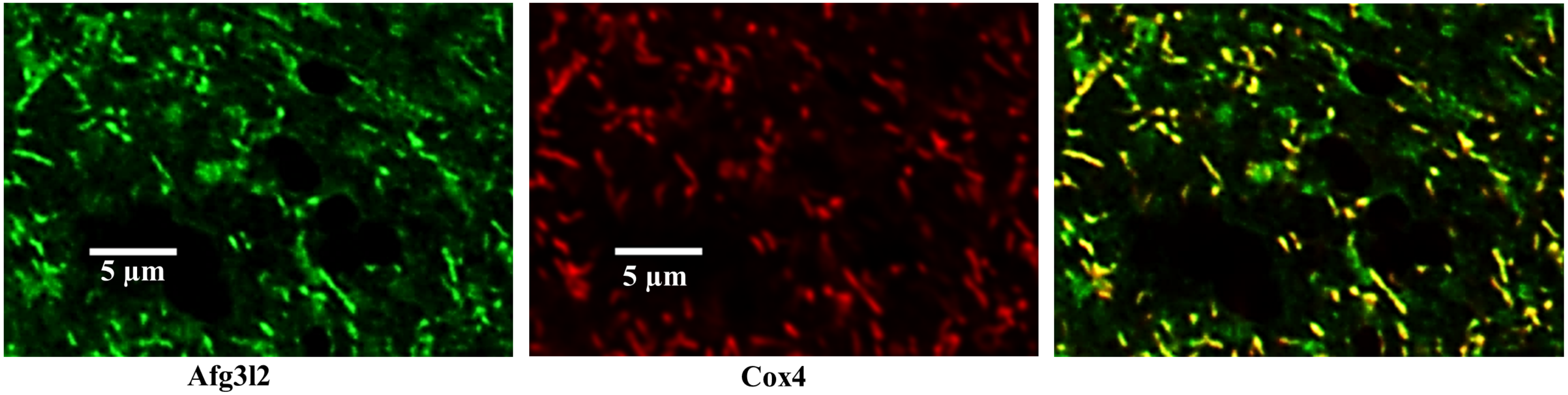


**b**

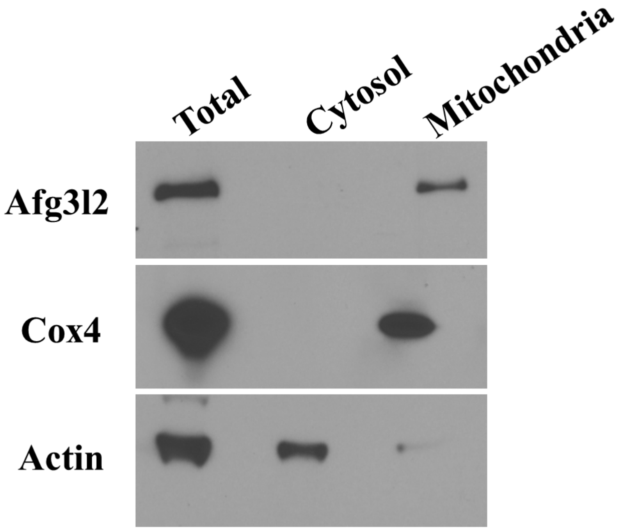

### Supplemental Figure 2

**a**

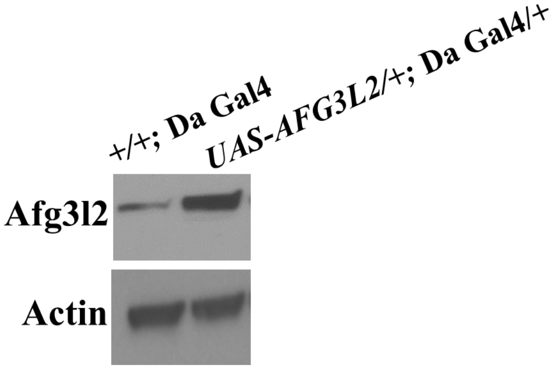


**b**

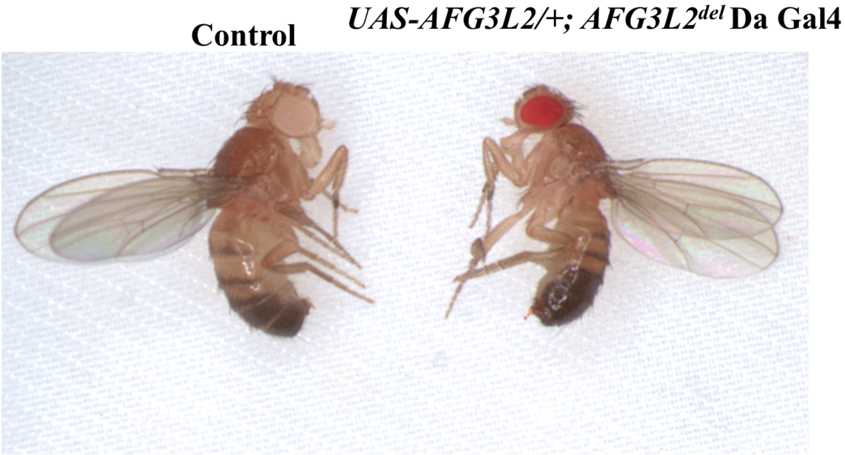

### Supplemental Figure 3

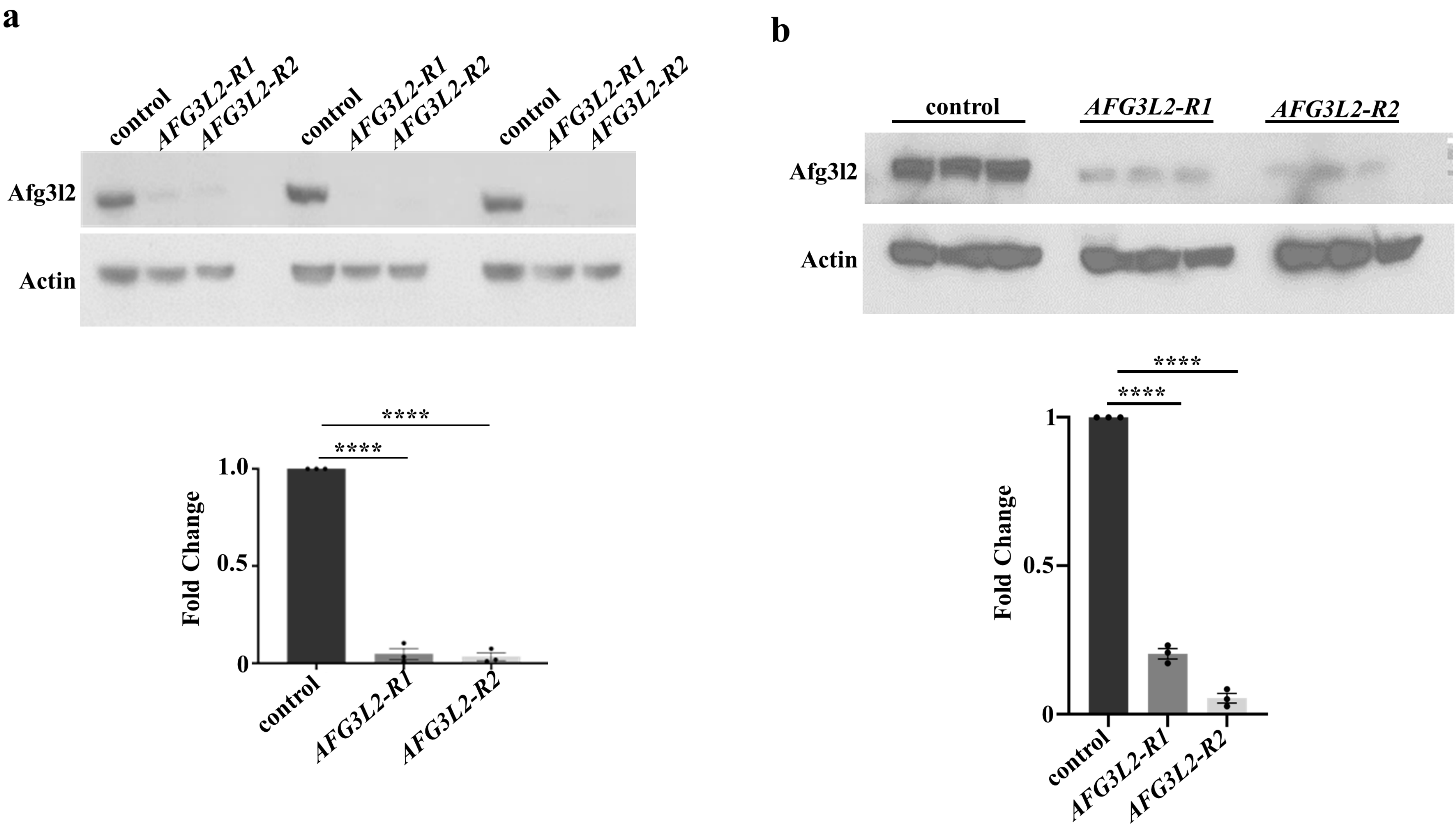

### Supplemental Figure 4

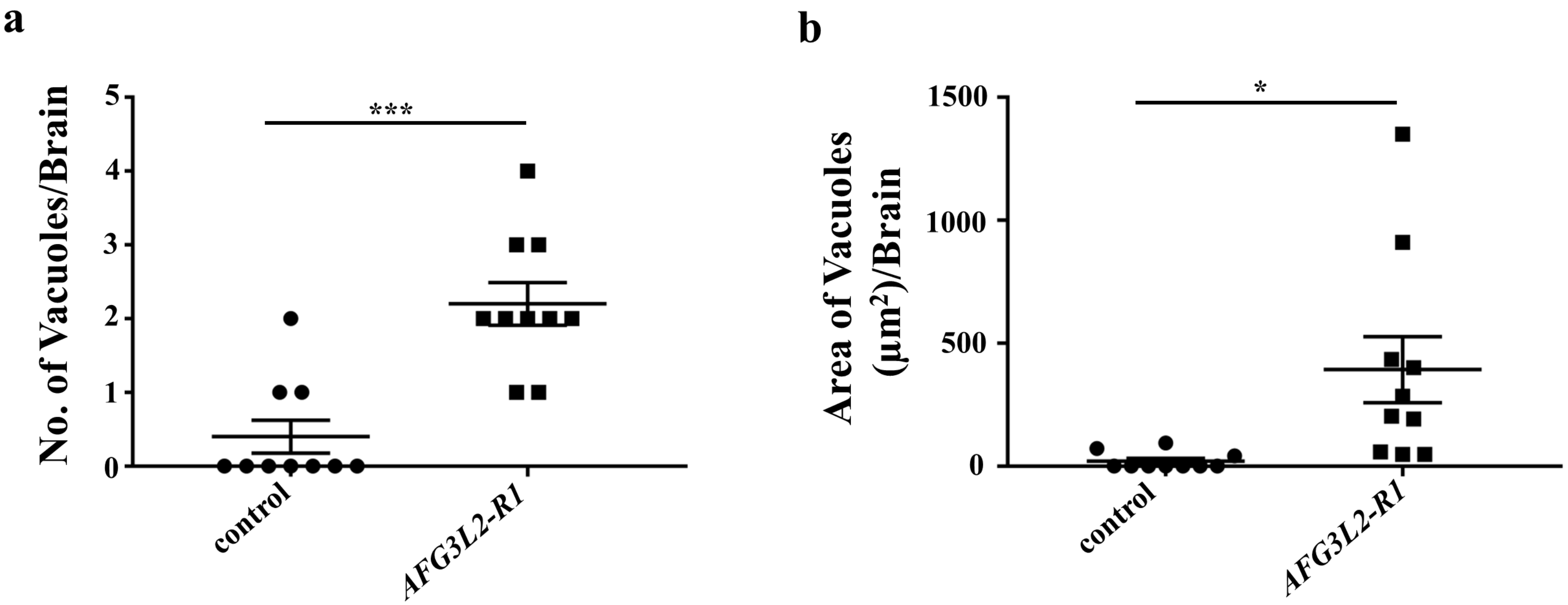

### Supplemental Figure 5

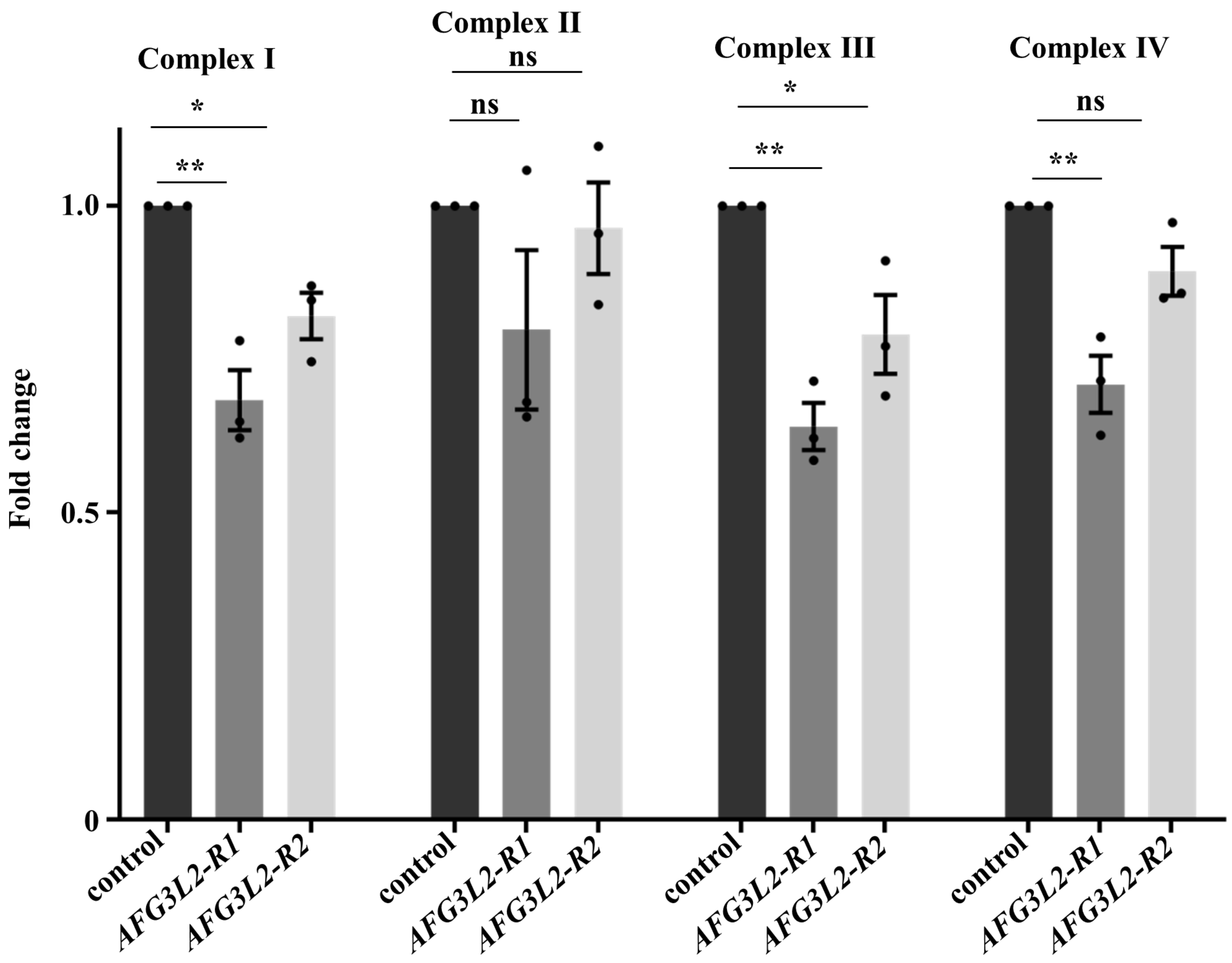

### Supplemental Figure 6

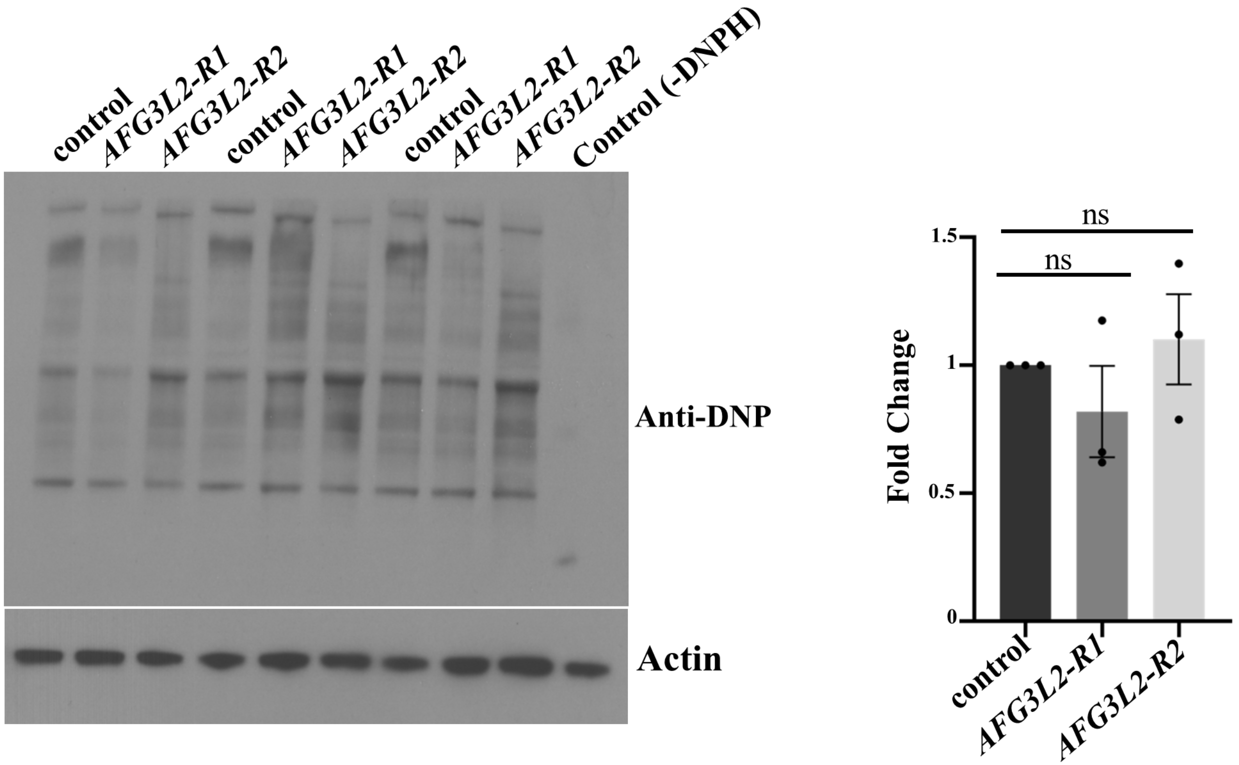

#### Supplemental Figure 7

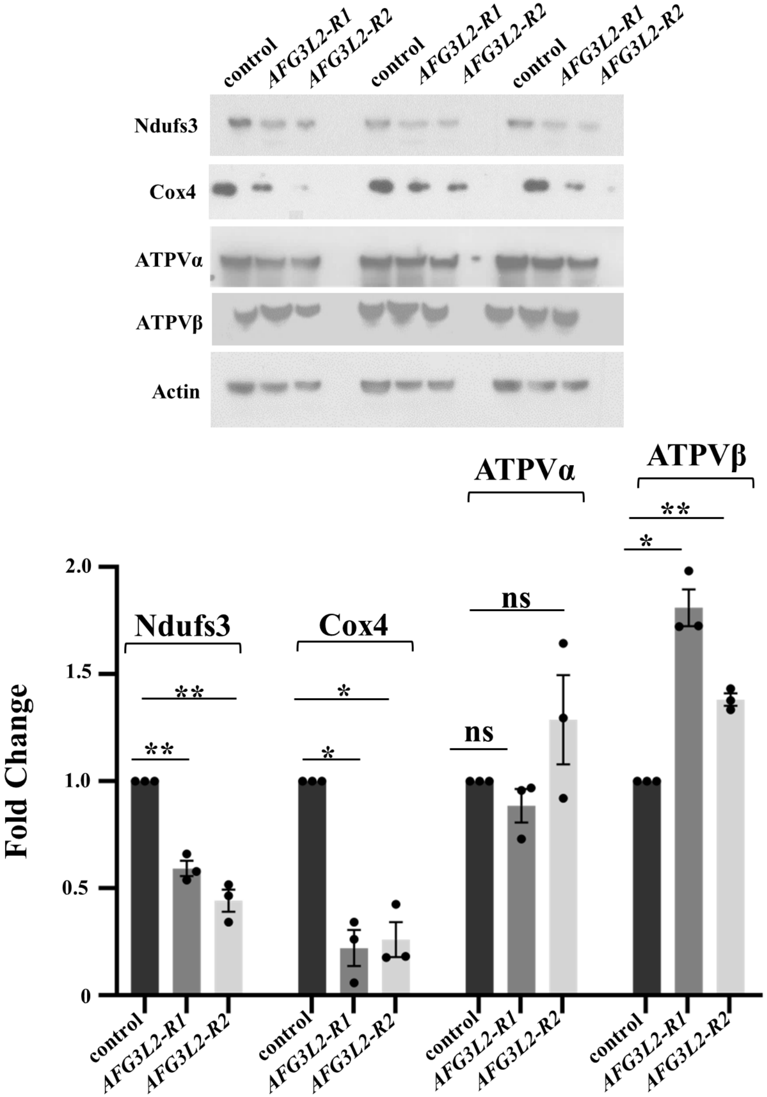

### Supplemental Figure 8

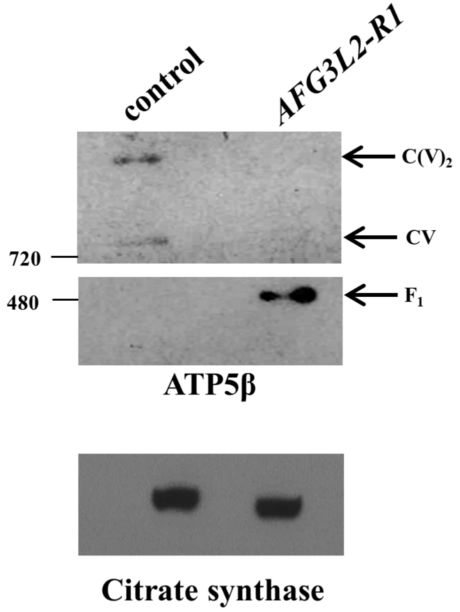

### Supplemental Figure 9

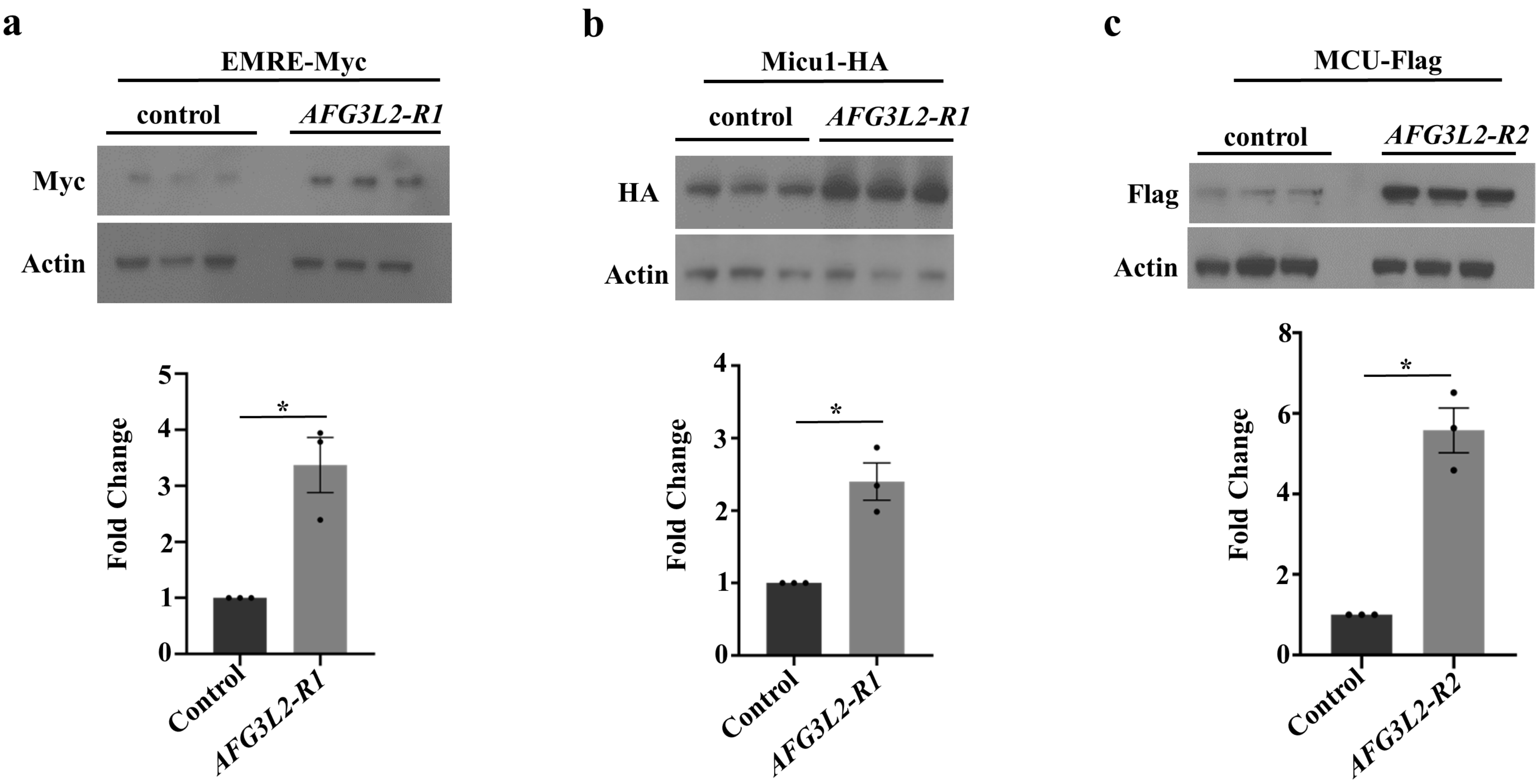

### Supplemental Figure 10

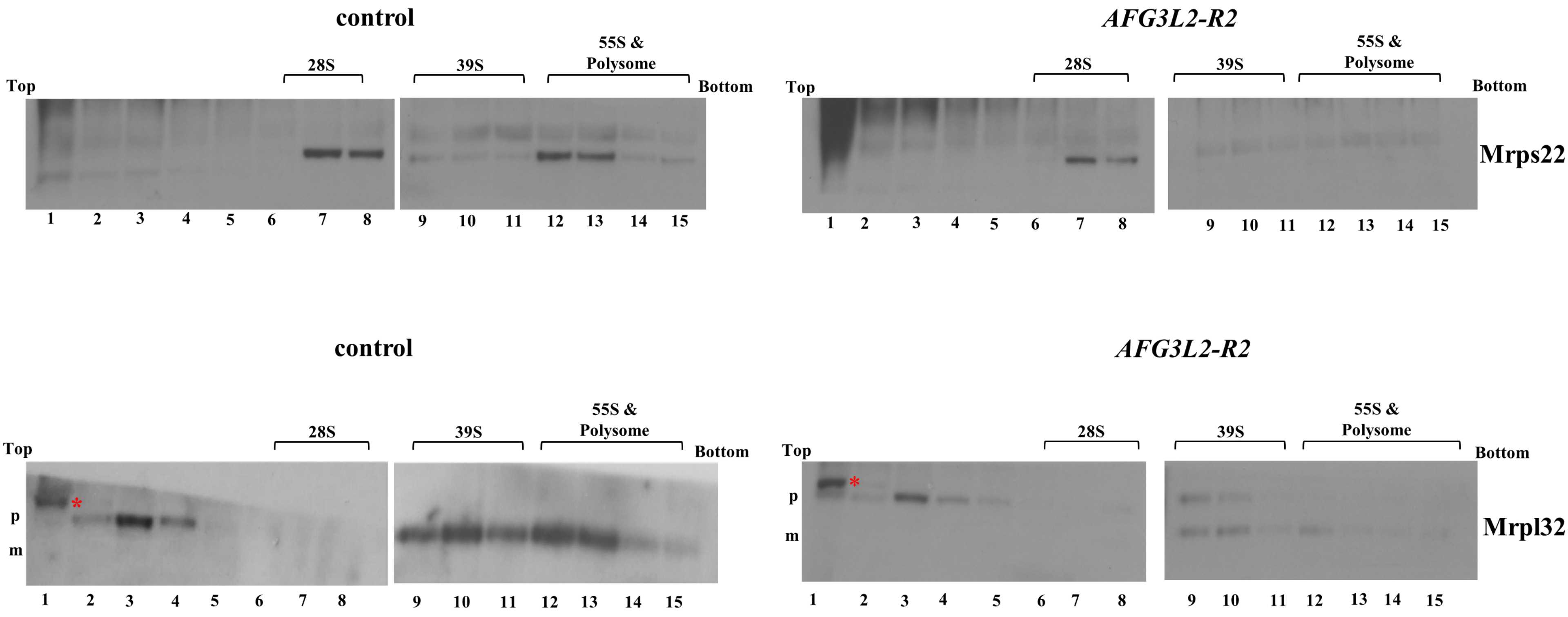

### Supplemental Figure 11

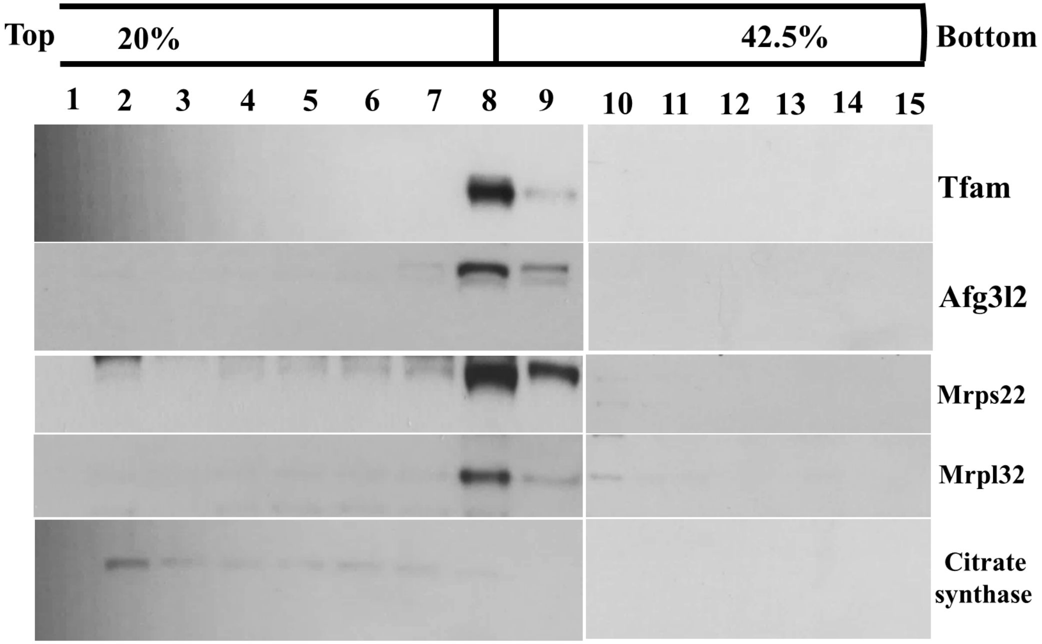

### Supplemental Figure 12

elav/CyoGal80; UAS-*LUCIFERASE* RNAi × UAS-*GFP or* UAS-*HSP60 or* UAS-*HSC70-5 or TRiP-OE LON;*UAS-*dCAS9-FLAG-VPR*

↓

elav/+; UAS-*LUCIFERASE* RNAi/UAS-*GFP*

elav/+; UAS-*LUCIFERASE* RNAi/UAS-*HSP60*

elav/+; UAS-*LUCIFERASE* RNAi/UAS-*HSC70-5*

elav/*TRiP-OE LON*; UAS-*LUCIFERASE* RNAi/UAS-*dCAS9-FLAG-VPR*

elav/CyoGal80; UAS-*AFG3L2-R2* RNAi × UAS-*GFP or* UAS-*HSP60 or* UAS-*HSC70-5 or TRiP-OE LON;*UAS-*dCAS9-FLAG-VPR*

↓

elav/+; UAS-*AFG3L2-R2* RNAi/UAS-*GFP*

elav/+; UAS-*AFG3L2-R2* RNAi/UAS-*HSP60*

elav/+; UAS-*AFG3L2-R2* RNAi/UAS-*HSC70-5*

elav/*TRiP-OE LON*; UAS-*AFG3L2-R2* RNAi/UAS-*dCAS9-FLAG-VPR*

**Supplemental Table 1**

| Genotype | Total Number (N) | Median Life Span (in days) |
|---|---|---|
| Elav (X) > UAS-*LUCIFEARSE* RNAi | 238 | 67 |
| Elav (X) > UAS-*AFG3L2-R1* RNAi | 465 | 9 |
| *w*$^{1118}$ | 480 | 66 |
| *AFG3L2*$^{del}$/+ | 548 | 64 |
| *SPG7*$^{del}$ | 196 | 34 |
| *SPG7*$^{del}$ ; +/+ ; *AFG3L2*$^{del}$/+ | 291 | 36 |
| Elav (II)> *UAS-GFP ;* UAS-*LUCIFEARSE* RNAi | 185 | 63 |
| Elav (II)> *UAS-GFP ;* UAS-*AFG3L2-R2* RNAi | 218 | 45 |
| Elav (II)> UAS-*AFG3L2-R2* RNAi/UAS-*HSP60* | 224 | 44 |
| Elav (II)> UAS-*AFG3L2-R2* RNAi/UAS-*HSC70-5* | 237 | 52 |
| Elav (II)> *TRiPOE LON*; UAS-*AFG3L2-R2* RNAi/ UAS-*dCAS9-FLAG-VPR* | 239 | 51 |

### Supplemental Table 2

| Gene name | Primer Sequence |
| --- | --- |
| RAP2L FORWARD | CCGCTGAAGGTAATGCCTTG |
| RAP2L REVERSE | CGTTTATCCGATCCTTTGCAGA |
| mt:tRNA:Lys-CTT FORWARD | GATGACTGAAAGCAAGTACTGG |
| mt:tRNA:Lys-CTT REVERSE | TCATTAGAAGTAAGTGCTAATTTAC |
